# An analysis of genetic diversity actions, indicators and targets in 114 National Reports to the Convention on Biological Diversity

**DOI:** 10.1101/2020.08.28.254672

**Authors:** Sean Hoban, Catriona Campbell, Jessica da Silva, Robert Ekblom, W Chris Funk, Brittany Garner, Jose A Godoy, Francine Kershaw, Anna MacDonald, Joachim Mergeay, Melissa Minter, David O'Brien, Ivan Paz-Vinas, Sarah Kim Pearson, Silvia Perez-Espona, Kevin Potter, Isa-Rita Russo, Gernot Segelbacher, Cristiano Vernesi, Margaret E Hunter

**Author notes:** Authors names are listed in the Cover Page to maintain double blind peer review.

## Abstract

Genetic diversity is critically important for all species-domesticated and wild- to adapt to environmental change, and for ecosystem resilience to extreme events. International agreements such as the Convention on Biological Diversity (CBD) have committed to conserve and sustainably and equitably use all levels of biodiversity-genes, species and ecosystems-globally. However, assessment and monitoring of genetic diversity are often overlooked, and there are large knowledge and policy gaps regarding genetic diversity conservation. In this study, we present the first quantitative analysis of genetic diversity assessments conducted by Parties to the CBD. We conducted a detailed, systematic analysis of 114 CBD 5th (submitted 2014) and 6th (submitted 2018) National Reports to quantitatively assess actions, progress on targets, values and indicators related to genetic diversity. First, we found that the importance of genetic diversity is recognised by most Parties to the CBD, and that recognition increased over time. However, genetic targets mainly addressed genetic diversity within cultivated plants, farm animals, and crop wild relatives, with little focus on other wild species. Also, actions for conserving genetic diversity primarily concerned *ex-situ* facilities and policy, rather than monitoring and intervention for maintaining genetic diversity *in situ*. The most commonly used indicators of genetic diversity status were the number of genetic resources in conservation facilities, number of threatened breeds, and Red List Index, which are not well correlated to genetic erosion in most species -- highlighting that genetic change is poorly monitored by current indicators. Lastly, analyses of genetic data observations, indigenous use and knowledge of genetic diversity, and strategies being developed and implemented to conserve genetic diversity are highly under-reported. We make several recommendations for the post-2020 CBD Biodiversity Framework to improve awareness, assessment, and monitoring, and facilitate consistent and complete reporting of progress of genetic diversity in future National Reports.

**Article Impact Statement:** An analysis of genetic diversity in CBD National Reports neglects non-domesticated species and demonstrates need for sufficient indicators.

## Introduction

Actions to halt the biodiversity crisis are increasingly urgent as human activities threaten life-supporting ecosystems and natural resources (Galli et al. 2014). In response, most countries have signed international accords, such as the Convention on Biological Diversity (CBD), committing to taking action and regularly reporting on progress towards protecting biodiversity. Although biodiversity includes diversity at the level of ecosystems, species, and genetics, the loss of genetic diversity (genetic and trait differences among individuals and populations within a species) has been relatively underappreciated for decades in both policy and practice, despite its importance (Vernesi et al. 2008, Laikre 2010, Holderegger et al. 2019). Genetic diversity provides wild species with the potential to adapt to environmental change (Wernberg et al. 2018), reduces negative inbreeding effects, supports ecosystem structure, integrity and resilience (Lotze et al. 2011), and is the basis for species diversity. It provides society with a range of options for plant and animal breeding to improve agriculture (Bhandari et al. 2017), forestry (Potter et al. 2017), fisheries (Houston et al. 2020) and other biodiversity-dependent industries, and forms the foundation for all other levels of variation (Hughes et al. 2008). Maintaining genetic diversity in domesticated species (e.g. breeds, landraces, and varieties) allows for cultivation/breeding under different environmental conditions and pressures such as pests and disease, which impact productivity (Hoffmann 2010).

Recent analyses show that genetic diversity has declined over the past century (Leigh et al. 2019), that genetically distinct populations are being lost, and that remaining genetic diversity is not well safeguarded *in situ* or *ex situ* (Khoury et al. 2019). Major drivers of genetic diversity loss include climate change, habitat fragmentation and destruction, overharvest, and small population sizes (CBD 2014). In spite of this, biodiversity assessments and reports tend to focus on species-level diversity. Within-species genetic variation assessments are limited (with some exceptions, see Hoban et al 2020; Santamaria and Mendez 2012) due to perceived costs and lack of training and standard protocols.

The CBD was first put into force in 1992 following the Rio de Janeiro Earth Summit and is the premier instrument for guiding and measuring global biodiversity conservation, sustainable development, and equity. The CBD’s signatory Parties (countries plus the European Union) committed to conserving all levels of biodiversity via 21 targets to be achieved by 2010 (CBD 2004), and a new set of 20 targets by 2020 (CBD 2010). However, the wording of targets emphasized genetic diversity primarily for species of direct human use, especially agricultural species. The 2010 genetic Target 3 focused on “crops, livestock, and harvested species of trees, fish and wildlife and other valuable species” and the 2020 Target 13 focused on “cultivated plants and farmed and domesticated animals and wild relatives, including other socio-economically and culturally valuable species” (CBD 2004; CBD 2010). Other commitments such as the Global Strategy for Plant Conservation (GSPC) and UN Sustainable Development Goals primarily relate to genetic diversity within agricultural species. The focus of targets likely influences how countries monitor, manage, and report genetic diversity information. Although genetic diversity is critical to agricultural innovations, resilience and food security, a primary focus on domesticated species could result in genetic erosion in other species (semi-managed and wild species, including wild relatives of domesticated species) and consequent losses in a range of ecosystem services. CBD is in the process of setting post-2020 goals, targets and indicators; however recent drafts frameworks still reflect lack of emphasis, ambition and clarity on genetic diversity conservation (Hoban et al. 2020).

Since 2000, signatory Parties have been required to submit National Reports every four years to report on their progress towards the CBD targets and on any related national targets that are implemented. These reports assist the Conference of Parties to consider lessons learned during implementation of the Convention, identify gaps in capacity and analysis at different levels (national, regional, and global), and formulate appropriate requests and guidance to signatory Parties and related bodies. CBD guidance differed slightly for each reporting period (see Methods and Discussion). Reports are typically compiled by government agencies, such as Ministries of the Environment, and other relevant stakeholders. In addition to reporting progress towards CBD and national targets, reports summarize biodiversity status, threats, advances in sustainable development, and inclusion of indigenous and local communities. Because of their periodic nature, global scope, and common template and guidelines for preparing the document, analyses of National Reports can help evaluate global progress in biodiversity conservation. Previously, National Reports have been analysed to assess: progress towards the implementation of the CBD targets (CBD 2014; Birdlife et al. 2016); national challenges in meeting CBD goals (Chandra & Idrisova 2011); indicators and knowledge gaps towards their use to achieve CBD targets (Bhatt et al 2019); gender equality (GmbH 2018); success on the implementation of the GSPC (Paton & Lughada 2011); and progress towards protected area management effectiveness targets (Coad et al. 2013). Most recently, 5^th^ National Reports were critically assessed to evaluate the status of the natural world and actions needed to conserve biodiversity and ecosystem services (IPBES 2019).

The consideration of genetic diversity, genetic approaches, or progress towards genetic diversity targets in National Reports has not yet been systematically analyzed. In fact, genetic diversity has been noted as a principal data gap to assessing biodiversity progress (OECD 2019). In previous analyses of National Reports, genetic diversity was mentioned primarily for agricultural species for global food security (IPBES 2019; CBD 2014; CBD Secretariat 2007). In the context of Aichi Target 16 (benefits and sharing of genetic resources), data gaps to monitor genetic diversity were also mentioned (Aguilar-Støen & Dhillion 2003). The scant analysis of genetic diversity in reports may be due to challenges in using genetic information for measuring CBD target progress (Chandra & Idrisova 2011), and lack of metrics for monitoring genetic diversity (Bubb et al. 2011; Walpole et al. 2009; OECD 2019).

In this context, we systematically assessed the consideration of genetic diversity in a large representative sample of 5^th^ and 6^th^ CBD National Reports (circa 2014 and 2018, respectively), to better understand how countries are assessing and protecting genetic diversity. This analysis contributes to a general understanding of how genetic diversity is considered in global biodiversity assessments, following numerous calls to increase such consideration (Shafer et al. 2015; Taylor et al. 2017). It also provides a basis for improvements to future reporting. Our specific aims were to analyze 5^th^ and 6^th^ National Reports to:

- Assess targets pertaining to genetic diversity, and which indicators (high level measures) are used to assess status (present state) and trends (change) in biodiversity
- Quantify the reporting of genetic diversity actions (e.g., management interventions, policy, funding), threats (e.g., concerns or drivers of change such as habitat fragmentation), and values (e.g., utility or benefits)
- Quantify the frequency with which different types of species are mentioned in reference to genetic diversity
- Determine whether results change across time and across socio-economic categories.

## Methods

We reviewed pairs of 57 5^th^ National Reports (NRs) and 57 6^th^ NRs-9 Spanish, 10 French, and 38 English for each report, available prior to 1 July 2019 (Supporting Information) from the CBD Clearinghouse (chm.cbd.int/). We evaluated reports using a structured questionnaire composed of standardized questions (hereafter “questionnaire”) developed over several phases in 2019. We devised the set of questions based on the CBD instructions to Parties on report preparation. We limited our questions to thematic sections that were common between the 5^th^ and 6^th^ NRs (see Supporting Information for an explanation of how each question matches CBD instructions). Every question had a detailed set of instructions to ensure consistent interpretation and completion of the questionnaire among reviewers.

The final questionnaire had 13 questions to assess each report. Here, we focus on nine of these questions (Table 1). This questionnaire was filled out by experts (the authors of this manuscript) in applied conservation genetics (hereafter “reviewers”), following a protocol similar to Pierson et al. (2016), Bhatt et al. (2019), and Chandra and Idrisova (2011). Each reviewer evaluated six to eight reports split between 5^th^ and 6^th^ NRs. Reviewers were allocated reports from multiple continents and with differing levels of economic income (according to the International Monetary Fund, IMF; World Bank levels are quite similar). In addition to reading the report “cover to cover,” the reviewers also performed a keyword search, querying the document for 15 pre-determined genetic diversity-related keywords (Supporting Information). After reading, highlighting and taking notes on each report, the reviewer completed the questionnaire (via a Google Form and compiled in a .csv file).

**Table 1:**
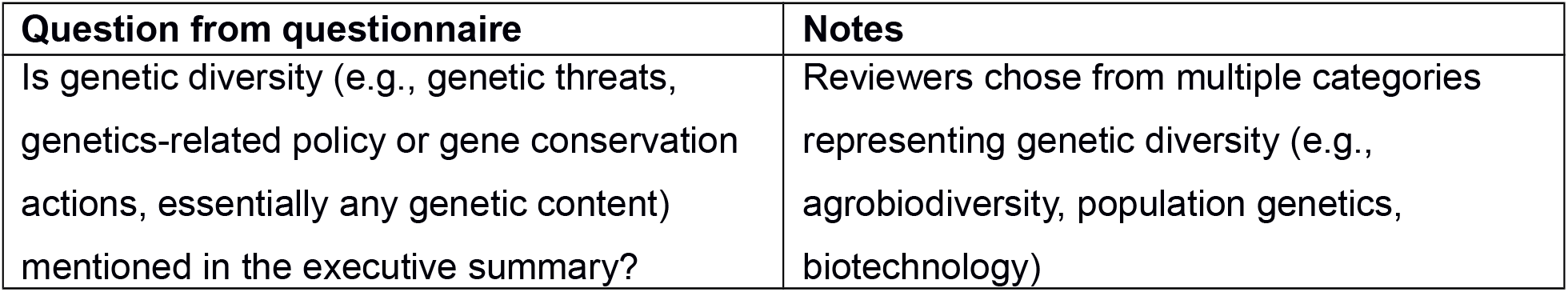
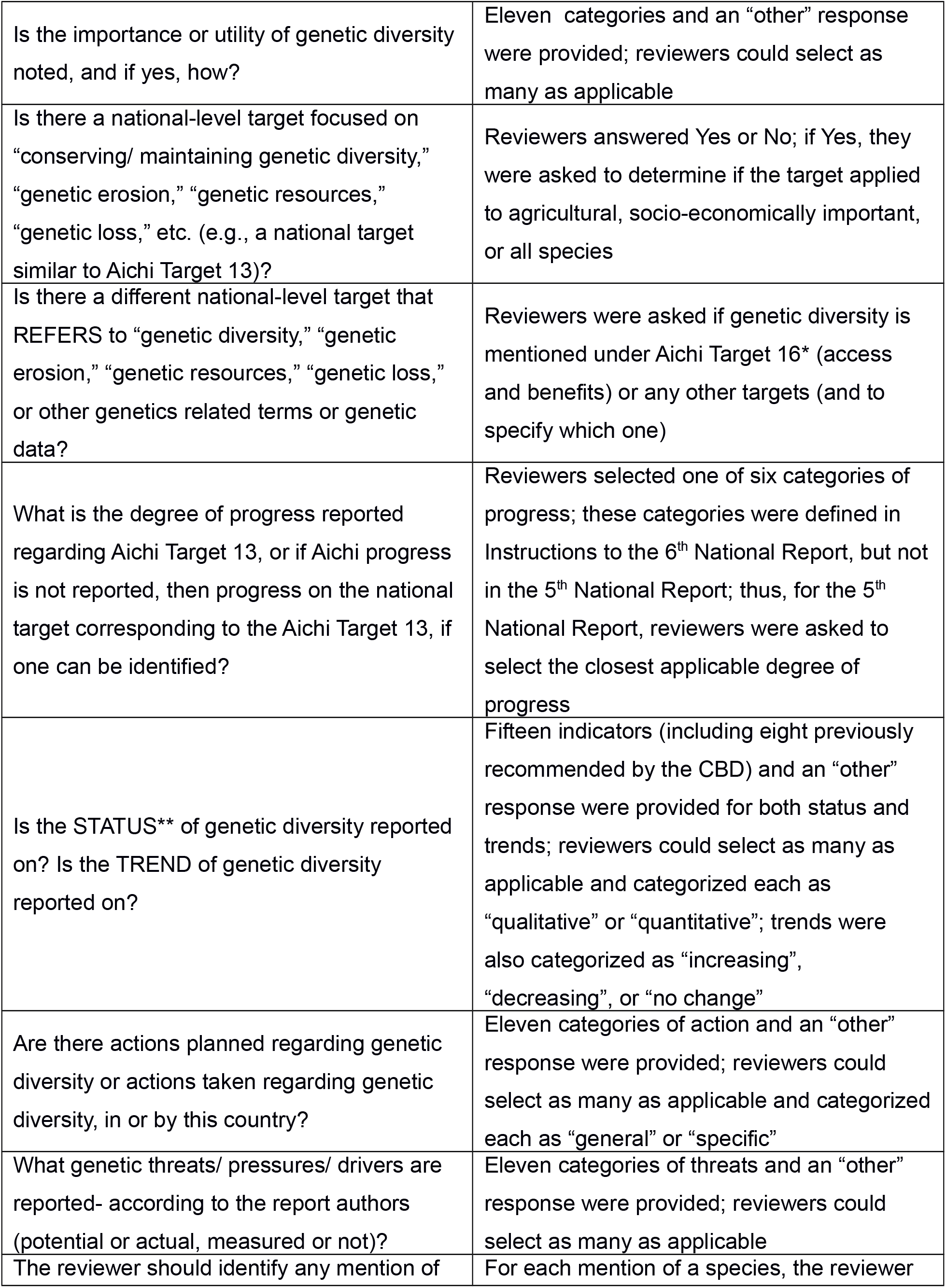
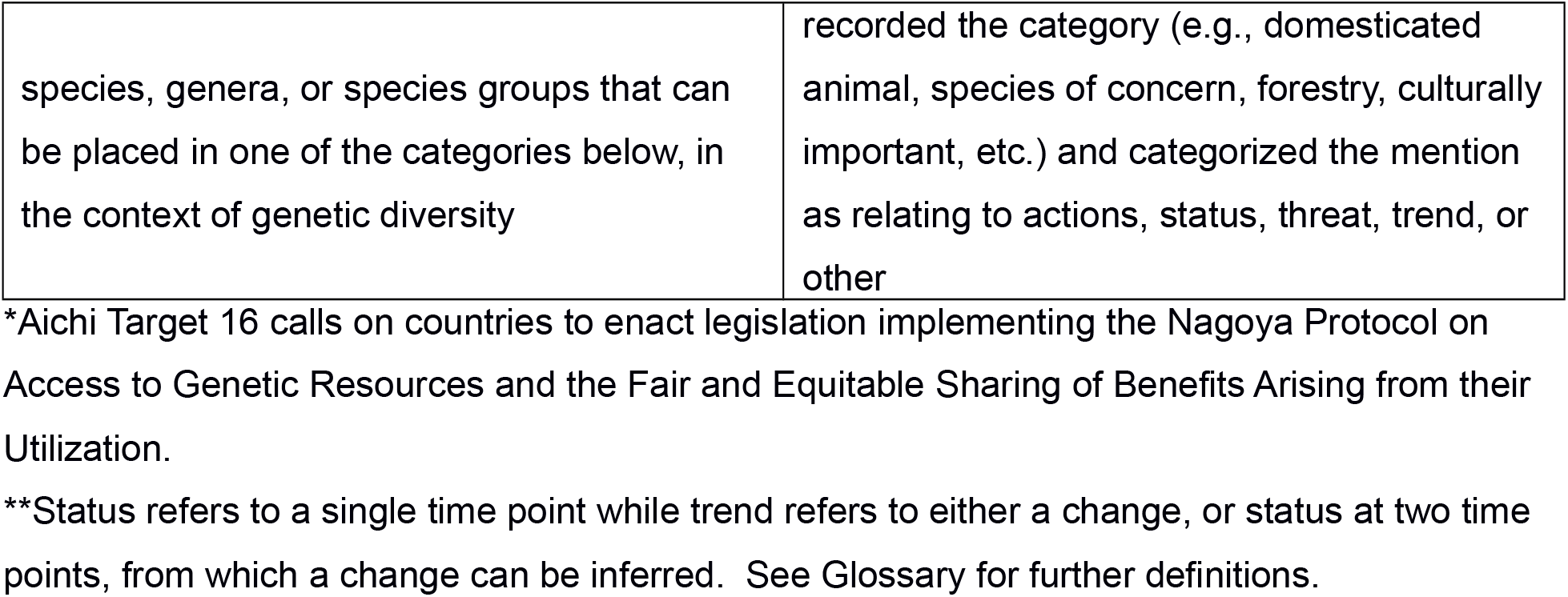
The questions answered by reviewers for this study.

We made a strong effort to standardize the review approach and to provide precise instructions and examples for reviewers. However, we acknowledge that each reviewer has a unique background, training, and expertise and there could be room for some level of interpretation in reviewing National Reports. To maximize standardization of questionnaire completion, we had several scoping phases to check agreement among reviewers and discussed disagreements as a group. Additionally, 15 reports (eight for 6^th^ and seven for 5^th^) were reviewed independently by two reviewers to ensure consistency in interpretation (Supporting Information).

Data were analyzed in R v3.6.3 (R Core Team 2020). To analyse the questionnaire responses, we first tabulated the number of reports with each potential answer. We compared the *proportion* of responses for each question between the 5^th^ and 6^th^ NRs using Fisher’s exact tests (for example, we determined whether the frequency of each category of indicators, e.g. the relative length of each bar in Fig. 1, differed between the reports). We also compared the mean *number* of responses recorded between the 5^th^ and 6^th^ NRs (for example the number of reports in which any genetic diversity indicators were identified, the sum of the lengths of bars in Fig. 1) using paired t-tests when data met the conditions of normality, and Wilcoxon tests when not. Responses from low-, medium-, and high-income countries (according to the IMF) were also compared for 5^th^ and 6^th^ NRs separately and pooled together.

**Fig. 1:**
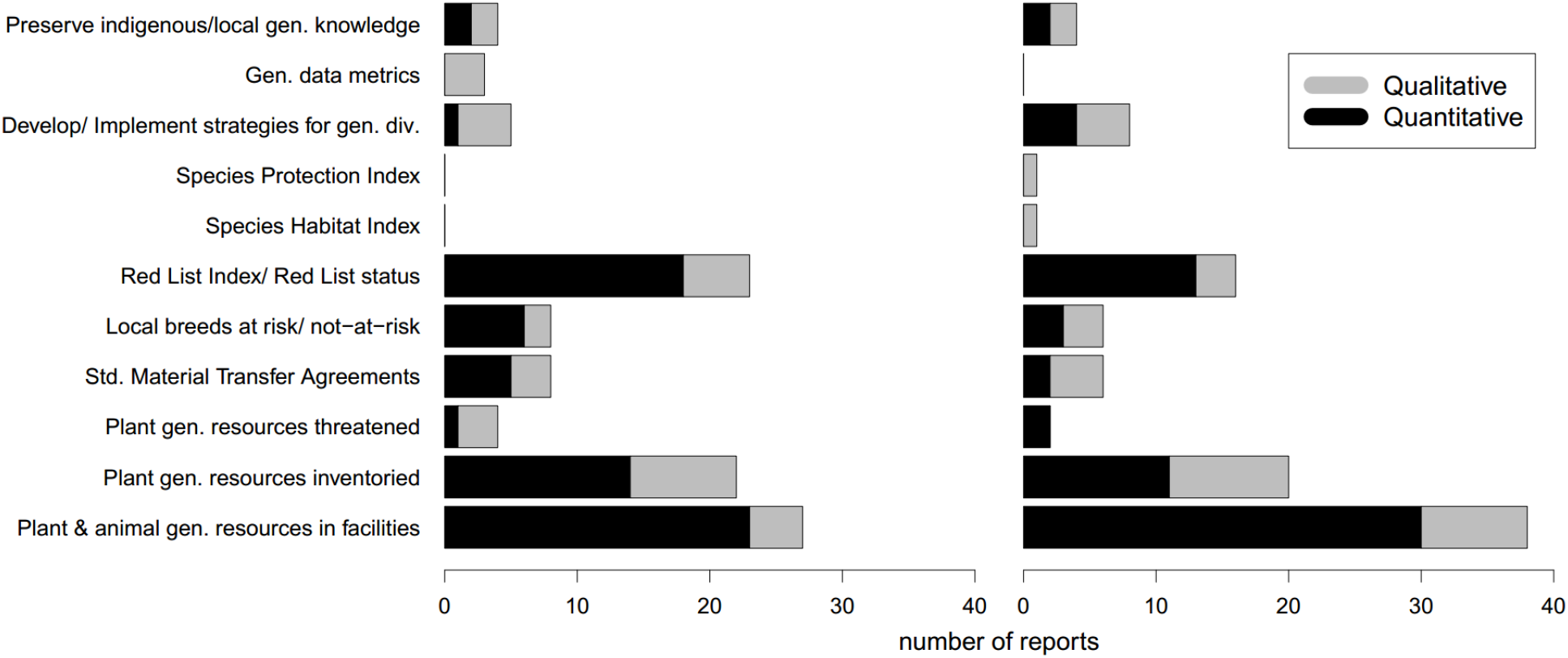
Number of 5th (left) and 6th (right) National Reports containing each indicator of genetic status with a Quantitative (numeric, such as a percentage) or Qualitative (descriptive, such as “high” or “low”) value.

## Results

Throughout the results section, differences between 5^th^ and 6^th^ National Reports (NRs) are only reported if statistically significantly different. Briefly, no contingency tests for comparing whether the categories changed between the 5^th^ and 6^th^ NRs were significant (i.e., no temporal change in frequency of responses in each answer category). The t-tests for the number of responses are noted if significant. All *p* values (significant and not significant) are reported in Supporting Information, Table S2. Results are almost identical when responses that were submitted as “Other” were recategorized into an existing response category (Supporting Information).

### Genetic diversity in executive summary

A majority (82%) of countries included an executive summary for the 5^th^ NR. Of these, 60% mentioned genetic keywords relating to agrobiodiversity (e.g., gene banks, breeds, or varieties); 40% mentioned genetic studies, gene conservation actions, or genetic processes; and 34% mentioned biotechnology or access to and benefit sharing of genetic resources (Supporting Information, Table S3).

### Values of genetic diversity

The most frequently noted values of genetic diversity included resilience to environmental or climate change (37% 5^th^ NR, 30% 6^th^ NR), increasing productivity in agriculture/ forestry/ fisheries (37% 5^th^ NR, 30% 6^th^ NR), developing new varieties in these sectors (26% 5^th^ NR, 25% 6^th^ NR), and adaptation to environmental change (26% 5^th^ NR, 23% 6^th^ NR, see Supporting Information, Table S4). There were 34% more mentions of values of genetic diversity in the 5^th^ National Report compared to the 6^th^ (*p*=0.04).

### Genetic diversity targets wording and progress

Of the 57 country reports reviewed, 70% and 79% referred to a national-level genetic diversity target in the 5^th^ and 6^th^ NRs, respectively (Supporting Information, Table S5). Many NRs (47% 5^th^ NR, 69% 6^th^ NR) mentioned other socio-economically important (e.g., not only agricultural) species relating to this genetic diversity target, while a smaller percentage included wording that could refer to species that do not have economic importance at present (21% 5^th^ NR, 38% 6^th^ NR). For target progress, the most commonly reported progress was “Some progress but insufficient” (57% 5^th^ NR, 44% 6^th^ NR), followed by “on track to achieve” (30% 5^th^ NR, 38% 6^th^ NR; Supporting Information, Table S6). Most countries also mentioned genetic diversity under Aichi Target 16, which regards access to and benefit-sharing of genetic resources (56% 5^th^ NR, 60% 6^th^ NR).

### Genetic diversity in other targets

This question assessed the extent to which countries identify genetic diversity as a concern, tool, or opportunity in association with any target other than Targets 13 and 16. The number of countries with references to genetics, associated with at least one Aichi target other than Targets 13 and 16, increased over time (19% 5^th^ NR, 49% 6^th^ NR), a significant increase of more than 2.5 times (*p*=0.001). In addition, the number of targets for which at least one country included a genetics-related reference increased over time. Genetics was mentioned under 13 different “other” Aichi targets in the 5^th^ NR, and under 19 “other” targets in the 6^th^ NR, a significant increase (*p*=0.044) of almost 50%. Several targets are of note. In the 6th NR, 16% of countries mentioned genetics in relation to both Targets 12 and 18, and 19% mentioned genetics for Target 19.

### Indicators used for genetic diversity status and trends

The most commonly mentioned indicators of status were the number of genetic resources in conservation facilities (Fig. 1, Supporting Information, Table S7), the number of plant genetic resources known/surveyed, and the Red List status. Meanwhile, the state of preservation of indigenous/local knowledge or use of genetic diversity was rarely reported, as were metrics from analysis of DNA/genetic markers.

Trends were mentioned half as often as status, and showed a mix of increasing, decreasing, and no change. A strong trend was only seen for “the genetic resources secured *ex situ*” which were typically reported as increasing Supporting Information, Table S8)

### Actions

The most common genetic diversity actions (Fig. 2) were establishing seed banks, research agencies or breeding programs, and laws or policies. Single time point genetic studies and genetic monitoring were rare.

**Fig. 2:**
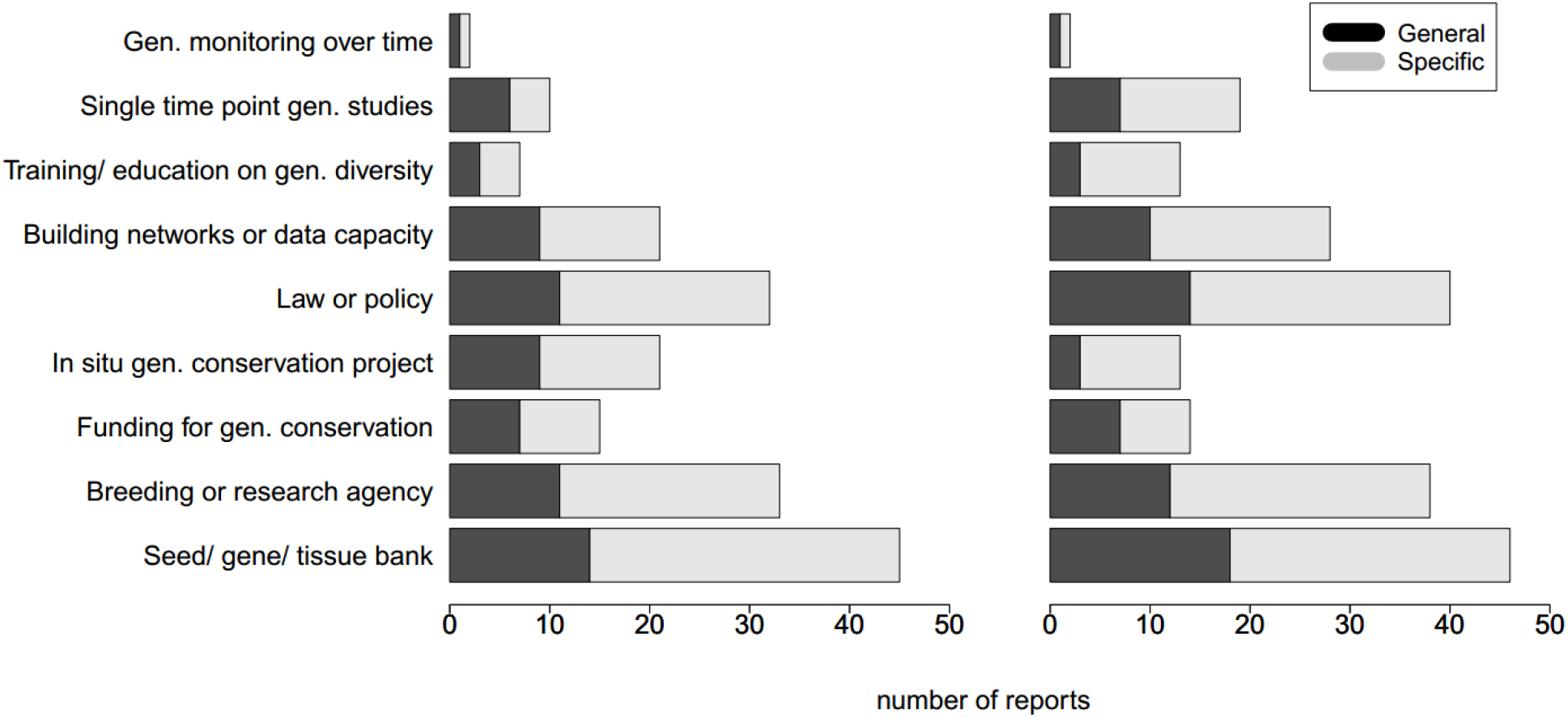
Number of 5th (left) and 6th (right) National Reports that report each type of genetic diversity action with General (e.g. non-specific action, black) or Specific (grey) information.

### Threats

National Reports documented a variety of recognized threats to genetic diversity. The most common threats in the NRs were replacement of native varieties or breeds, habitat fragmentation, and climate or environmental change (Supporting Information, Table S9). Other typical conservation genetic concerns were also mentioned, including decrease in range size, overharvest, pests or invasive species, small population problems, genetic modification, and hybridization. More threats were identified in the 5^th^ National Report than the 6^th^ (*p*=0.02).

### Species mentioned

The top species types (Fig. 3; Supporting Information, Table S11) mentioned were cultivated crops and farm animals (both >20% in both reports), followed by crop wild relatives, forestry species, and species of conservation concern (9 to 11% in both reports). There were few references to “other socio-economically important species” such as wild-harvested species, species providing ecosystem services, and culturally valuable species or wild relatives of domesticated animals (all <5% both reports). Species were most frequently acknowledged in relation to “actions,” followed by mentions relating to “threat” or “status” of genetic diversity (Fig. 3). References of species’ “change” in genetic diversity were rare.

**Fig. 3:**
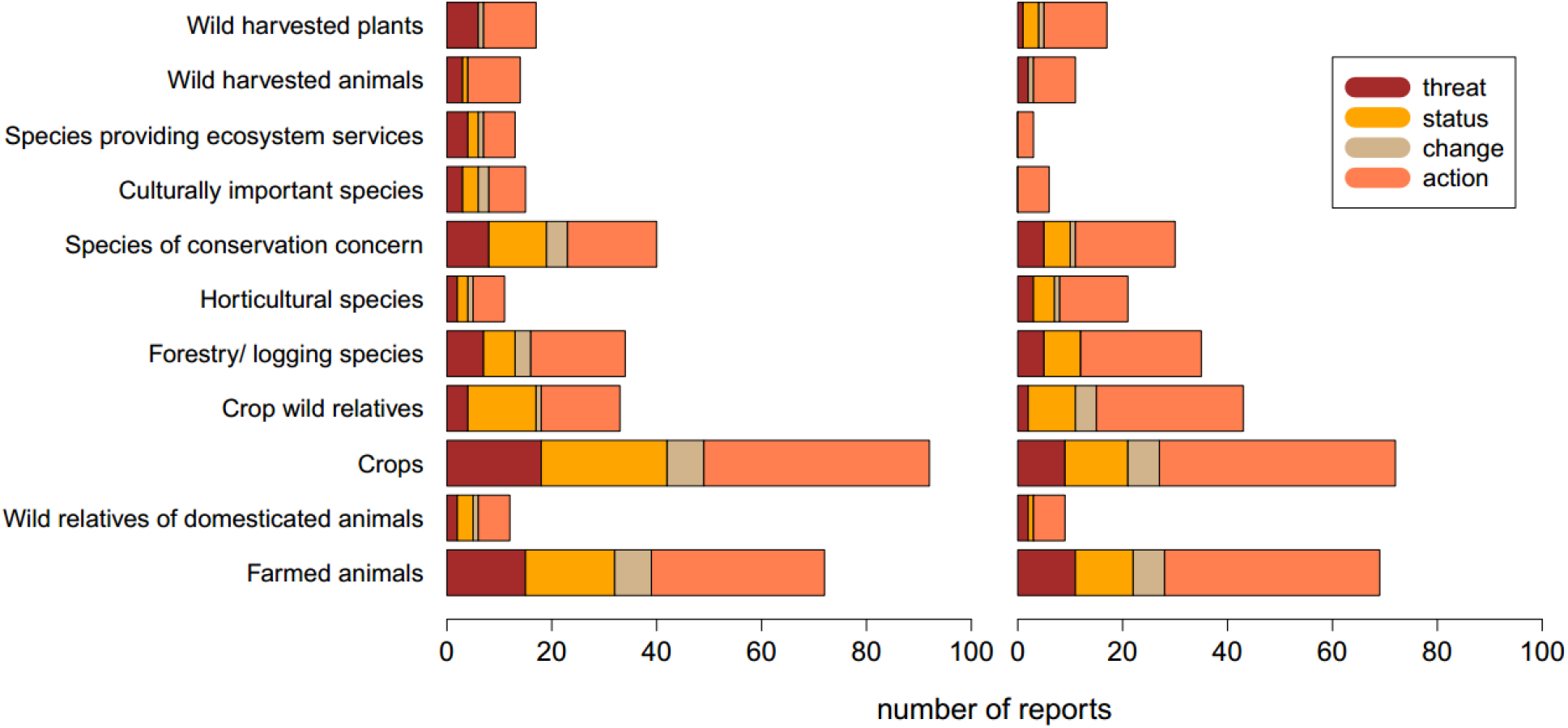
Number of mentions (x-axis) where each species type (y-axis) is mentioned in four different contexts (whether the mention of the species’ genetic diversity related to a threat, status, change, or action) for each report.

### Differences related to income levels of countries

Differences in countries’ responses according to their income level were only observed in three areas: mentions of genetic diversity in relation to targets other than Aichi Targets 13 and 16, threats, and species types.

Low-income and middle-income countries were more likely than high-income countries to mention genetics in relation to genetic diversity targets, genetic resources targets, and in targets other than Aichi Targets 13 and 16, though none of these results were significant (Supporting Information).

There were no significant differences among income level categories in the number of mentions of threats, but there were significant differences in the proportion of categories of threat. Specifically, NRs from middle- and low-income countries had fewer mentions of small population size and habitat fragmentation as threats to genetic diversity, but more mentions of replacement of traditional varieties as a threat (significant only when 5^th^ and 6^th^ NRs are pooled, *p*=0.04).

As for species type, NRs from middle- and low-income countries had fewer mentions of species of conservation concern and species providing ecosystem services, but had more mentions of horticultural species, compared to high-income countries (only marginally significant when 5^th^ and 6^th^ NRs are pooled, *p*=0.08).

## Discussion

Maintaining genetic diversity provides species with adaptive potential, contributes to resilient, functioning of ecosystems, and is necessary to counter current global crises (e.g., climate, biodiversity loss, hunger; Di Falco & Perrings 2003; Reusch et al 2005; Sjöqvist & Kremp 2016; Raffard et al. 2019). If countries are to effectively design and measure progress on targets and interventions for this goal, they need to adequately report on genetic diversity. The CBD National Reports are an essential information source, and reveal how countries value, monitor, and manage biodiversity. For 5th and 6th CBD National Reports we found that most countries mentioned the importance of genetic diversity. However, when status and actions relating to genetic diversity were reported, they primarily referred to agricultural species, and used indicators that are not well connected to genetic diversity status or change. There were only minor differences in findings for the 5th vs. 6th Report, reflecting consistency over time in spite of some differences in guidance on National Reports. Below, we discuss the findings and how they point to improved future National Reports and genetic diversity Targets, indicators and capacity.

### Genetic diversity awareness and targets

Encouragingly, most countries have sufficient awareness of genetic diversity to include it in the Executive Summary and to have a genetic diversity National Target. Numerous benefits of genetic diversity were recognised, including maintaining ecosystem stability or services, food production, and pest/disease resistance. National Targets largely, but not always, focused on agricultural species, which aligns with how Aichi Targets 13 and 16 approach genetic diversity. The high (and increasing over time) number of reports linking genetic diversity to Target 12 (prevention of species’ extinctions) may reflect increasing recognition of genetic diversity’s role in supporting species survival (Booy et al. 2000; Sgrò et al. 2011). Meanwhile, references to genetic diversity in Targets 18 (recognition, respect and integration of traditional knowledge for biodiversity conservation) and 19 (improving scientific knowledge of the values, functioning, and status of biodiversity, and the consequences of its loss) may reflect increasing recognition of the importance of indigenous and scientific knowledge on genetic diversity.

Low- and middle-income countries were more likely than high-income countries to discuss genetic diversity in relation to targets other than Aichi Targets 13 and 16. This may reflect higher interest in knowledge transfer and greater recognition of traditional ecological knowledge by these countries. Nonetheless, the importance of genetic diversity and the use of genetic approaches are still rarely mentioned. We emphasize a need for increased knowledge of, capacity development, and access to genetic data for practical use (Recommendations 1 and 2 below) and for policy makers (in CBD and beyond) to explicitly state that genetic diversity is important for multiple targets in reporting guidelines (Recommendation 6).

### Genetic diversity indicators

The number of genetic resources in conservation facilities, the number of plant genetic resources known/surveyed, and the Red List were commonly-mentioned indicators of the status of genetic diversity. These indicators are recommended by the CBD (Supporting Information), but their prevalence probably also reflects data availability in existing national and international databases. The first two indicators also likely reflect Aichi Target 13’s emphasis on agricultural species, and the importance of genetic resources for national food security (Esquinas-Alcázar 2005; Khoury et al. 2014). Despite their status as official genetic CBD indicators, they do not directly assess genetic diversity and are at best loosely connected to genetic change (Hoban et al. 2020). The CBD is not the only place in which imperfect substitutes for genetic diversity are used as Indicators. For example, the Montréal Process (an international forest sustainability framework) includes three “genetic diversity” indicators: species at risk of losing genetic variation; population levels of forest-associated species; and the status of on-site and off-site gene conservation efforts (montrealprocess.org). Overall, the paucity of reliable indicators of genetic diversity shows that the scientific community needs to develop affordable, standardized indicators that clearly track genetic change (Hoban et al. 2020; Laikre et al. 2020) (Recommendation 4).

Only 5% of countries referenced DNA-based studies (e.g. genetic statistics) or recorded indigenous and local knowledge or maintenance of genetic diversity. This is likely because there are no official CBD indicators based on genetic markers or indigenous knowledge or awareness of the importance of genetic diversity (see Recommendation 3). Further, indicator uptake and reporting depend on availability of data on baselines and trends, as well as capacity (resources, technical expertise, and management) (Vanhove et al. 2017). Similarly, only 13% of countries reported measures to develop or implement strategies for minimizing genetic erosion, even though Aichi Target 13 called for such strategies. One limiting factor may be that there is no available database on, or guidance on how to develop and apply, such strategies.

### Genetic diversity actions

The most common actions reported related to the conservation of genetic diversity were *ex situ* strategies (seed banks, research agencies, laws, etc.). Less commonly mentioned were *in situ* actions (developing *in situ* genetic conservation projects, single time point genetic studies). The lower frequency of reporting on these actions may reflect the constraints on capacity, knowledge, and funding constraints previously mentioned. While *ex situ* actions, laws and policies are important to forestall absolute loss of genetic diversity, they cannot be relied upon exclusively, because only a limited representation of species genetic diversity can be maintained *ex situ*, and sustaining genetic diversity *in situ* is important to allow for natural processes such as adaptation to environmental change (Sgrò et al. 2011). Further, *ex situ* actions do not address two other major reported threats (habitat fragmentation and climate or environmental change).

Genetic data (as indicators) and genetic monitoring (as an action) were mostly absent from reports, even though genetic monitoring programs exist (e.g. European trees: Aravanopoulos et al. 2015; California Chinook salmon: Meek et al. 2016; Swiss grey wolves: Dufresnes et al. 2019). These findings complement the observation by Pierson et al. (2016) that European countries rarely include genetic diversity data and monitoring in species recovery plans due to a lack of legislative requirements, and relatively limited involvement of geneticists (Taft et al. 2020). While one-time genetic studies are important as a first step, assessing and reporting on change in genetic diversity requires standardized genetic monitoring across long-term time frames (i.e., multi-decadal) (Hoban et al. 2014; Mathieu-Bégné et al. 2019). Although requiring significant resources, capacity, and strategic planning timeframes, long-term monitoring programs will improve understanding of genetic diversity status and trends and meet the goals of the CBD and other international commitments.

### Types of species

References to genetic diversity in National Reports were biased towards cultivated crops and farm animals. Genetic diversity of crop wild relatives, forestry species, and species of conservation concern was discussed to a lesser degree, while there were few mentions of “other socio-economically important species” such as wild harvested species, species providing ecosystem services, and culturally valuable species. Because trends in one of these groups does not necessarily correlate to change in other species groups, monitoring and reporting should be conducted on multiple representatives of each “group” (Hollingsworth et al 2020). Our observations may reflect the relative emphasis of the Aichi Target 13 wording, as well as a focus on economic values. We support calls for the CBD to explicitly state the importance of maintaining genetic diversity of all species (Laikre et al. 2020), as agriculturally valuable species make up a small fraction of life on Earth (Recommendation 3).

### Change over time

The relative *frequencies* in each category or response for all questions were not significantly different between 5^th^ and 6^th^ National Reports. For example, reporting on “seed/tissue/gene banks” was the most frequent action in both reports. This suggests that the focus and priorities of countries regarding genetic diversity have not shifted substantially over this four-year period. However, the 6^th^ National Report contained significantly more mentions of genetic diversity related topics under Aichi Targets not specifically focused on genetic diversity (i.e. Targets other than 13 and 16). Instructions did not ask countries to mention genetics with respect to other Targets in either report, so this increase may be attributable to increasing awareness and affordability of genetic approaches and knowledge of genetic diversity, and to numerous calls for consideration of genetics in biodiversity policy (Pierson et al. 2016; Shafer et al. 2015; Taylor et al. 2017). Second, significantly more trends, values, and threats to genetic diversity were identified in 5^th^ Reports. This could be because instructions for 5^th^ National Reports provided more detail about these aspects (Supporting Information). If this is true, it emphasizes that the wording of CBD instructions (not just Targets) is extremely important for reporting (Recommendations 3, 5 and 6). We also note that Trends were reported less than half as often as Status: National Reports seriously lack temporal comparison of progress of biodiversity conservation over time (see also Bhat et al 2020), emphasizing a need for monitoring (Recommendations 4 and 6).

### Caveats

We acknowledge several methodological caveats (expanded upon in Supporting Information). Although the CBD intended consistency across National Reports to facilitate tracking progress (cbd.int/reports/guidelines/), the structure and formats were somewhat different. Therefore, in our study we focused on equivalent sections between the reports: status, threats, actions, obstacles, and progress towards the different biodiversity conservation targets. We also note that National Reports do not reflect all conservation actions or knowledge in a country, but are summaries of activities and progress, and might represent a country’s priorities, as well as data availability. Report writers may lack sufficient access to data, knowledge of actions, or expertise when preparing the reports. Thus, there may be discrepancies between reported progress and actions, and actual progress within countries (i.e., some countries may have achieved more or less than they reported, see Recommendation 6). However, there is no other global database on how countries assess their genetic diversity. Despite these caveats, we were able to analyze data provided in National Reports and discuss patterns and trends in reporting.

### Recommendations and Conclusions

Here, we make recommendations for addressing issues we uncovered in our detailed survey of 114 National Reports. First, our assessment suggests general work needed to improve measures for assessing genetic diversity in the post-2020 world.

1. **Increase awareness and knowledge sharing of the essential role of genetic diversity in biodiversity, and the benefits it brings to nature and people**. Assessments of genetic diversity in agricultural and natural systems must be communicated to policy makers and conservation managers via networking and capacity building, and the effectiveness of such science-policy communication should be measured (e.g. Lundmark et al. 2019).
2. **Increase quality and quantity of reporting on genetic diversity in wild species, especially long-term monitoring to evaluate trends**. This must be made feasible for all countries, not simply those with highly-developed scientific infrastructure. Collaborations between geneticists and conservation managers can provide guidance on cost-effective monitoring (Perez-Espona et al. 2017). We also provide recommendations for CBD reporting post 2020. Signatory countries and the CBD Secretariat invest significant resources in designing, producing, and showcasing reports, so it is beneficial for reports to be informative and enable comparisons of progress over time.
3. **Reporting on the genetic diversity in all species, not just species associated with agriculture, is essential to maintain stable, resilient ecosystems**. We recommend that the CBD request reporting of targets, status, threats, and actions relating to genetic diversity in each category of species (Fig. 3) and in major taxonomic groups.
4. **The CBD should consider recent suggestions on genetic goals, action targets, and genetic diversity indicators** (Hoban et al. 2020) which are easy to quantify, reliably reflect changes in genetic diversity and enable evaluation of actions taken to protect and restore genetic diversity. Indicators and data relating to indigenous and local use and knowledge of genetic diversity are also needed.
5. **The CBD should adopt improved and consistent terminology relating to genetic diversity and genetic approaches**. Clear definitions of terms in reporting instructions via glossaries, with examples, at least in the six major CBD languages should improve consistency in reporting, and awareness of the importance of genetic diversity conservation.
6. **Enable clearer comparisons of reporting on genetic diversity among countries and over time**. We suggest clearer structure for National Reports and greater guidance to countries on what to report and how. As one example, National Report formatting could include lists of actions performed, or categories of species managed, which can be “checked off”. Meanwhile, to allow unstructured information, the CBD could encourage inclusion of additional reports and/or case studies, e.g. Hollingsworth et al 2020.

Implementing these recommendations should increase reporting of indicators, actions, and progress towards the protection of genetic diversity in all species, as well as consideration of genetic diversity in other targets. They also represent an opportunity for CBD to facilitate capacity building and international collaboration, to equip countries to monitor genetic diversity post-2020. Of course, conservation geneticists need to actively participate at the science-policy interface, and we anticipate their increased inclusion to progress discussions of CBD and global work on genetic conservation.

## Supporting Information

Supplemental Methods, Results and Discussion (Appendix S1), and CBD Instructions (S2), and Questionnaire Details (S3, S4 and S5) are available online. The authors are solely responsible for the content of these materials. Queries should be directed to the corresponding author.

## Glossary

Action: an activity undertaken (or planned to be undertaken) by a country to make progress towards one or more targets (e.g. development of policy; management intervention; training; implementation of a conservation program)
Aichi targets: a set of 20 targets agreed by the CBD to be achieved by 2020
CBD: The Convention on Biological Diversity
Genetic diversity: inherited genetic and trait differences that vary among individuals and populations within a species
Genetic erosion: a loss of genetic diversity
Genetic resource: genetic material of actual or potential value. Genetic material is any material of plant, animal, microbial or other origin containing functional units of heredity (CBD, Art 2, see also https://biodiversity.europa.eu/topics/genetic-resources). Often used to refer to species diversity, e.g. number of plant wild relative species
Indicator: a measure used to present a high level summary of biodiversity; we include in our questionnaire official CBD indicators and other indicators
National Reports: reports submitted by signatories (countries) to the CBD every 4 years to outline progress towards CBD and national targets: 5th Reports were submitted starting in 2014 and 6th Reports were submitted starting in 2018
National targets: targets that each country sets for themselves: a national-level interpretation of the 20 CBD Aichi targets
Progress: an assessment of whether a country considers itself as on track to meet a CBD or national target, for example: “on track to achieve”; “some progress but insufficient”; “moving away”
Status: a measure of genetic diversity (or more frequently a proxy assumed to relate to it) at a single time point, e.g., the number of seeds in a seed bank at a given point in time.
Threat: a process or driver of change that is, or has potential to be, detrimental to genetic diversity
Trend: a measure of change in status over a period of time, i.e. an observation that status has increased, decreased, or has not changed.
Value: a perceived utility or benefit from genetic diversity

## References

1. Aguilar-Støen M, Dhillion SS. 2003. Implementation of the Convention on Biological Diversity in Mesoamerica: environmental and developmental perspectives. Environmental Conservation 30: 131–138. https://doi.org/10.1017/S0376892903000110

2. Aravanopoulos FA, et al. 2015. Development of genetic monitoring methods for genetic conservation units of forest trees in Europe. European Forest Genetic Resources Programme (EUFORGEN), Bioversity International, Rome.

3. Bhandari HR, Bhanu AN, Srivastava K, Singh MN, Shreya HA. 2017. Assessment of genetic diversity in crop plants - an overview. Advances in Plants & Agricultural Research 7: 279–286. https://doi.org/10.15406/apar.2017.07.00255

4. Bhatt R, Gill M, Hamilton H, Han X, Linden H, Young B. 2019. Uneven use of biodiversity indicators in fifth national reports to the Convention on Biological Diversity. Environmental Conservation 1–7. https://doi.org/10.1017/S0376892919000365

5. Birdlife International, et al. 2016. Score Card - Convention of Biological Diversity: Progress Report towards The Aichi Biodiversity Targets. https://www.birdlife.org/sites/default/files/score_card_booklet_final.pdf (accessed 18 August 2020).

6. Booy G, Hendriks RJJ, Smulders MJM, Groenendael JM, Vosman B. 2000. Genetic Diversity and the Survival of Populations. Plant Biology 2: 379–395. https://doi.org/10.1055/s-2000-5958

7. Bubb P, et al. 2011. National indicators, monitoring and reporting for the strategic plan for biodiversity 2011-2020. A review of experience and recommendations in support of the CBD. Report UNEP/CBD/AHTEG-SP-Ind/1/INF/2. https://www.cbd.int/doc/meetings/ind/ahteg-sp-ind-01/information/ahteg-sp-ind-01-inf-02-en.pdf (accessed 13 July 2020).

8. CBD. 2004. Decision VII/30. Strategic Plan: future evaluation of progress. https://www.cbd.int/doc/decisions/cop-07/cop-07-dec-30-en.pdf (accessed 13 July 2020).

9. CBD. 2010. Decision X/2. Strategic Plan for Biodiversity 2011-2020. Available from https://www.cbd.int/doc/decisions/cop-10/cop-10-dec-02-en.pdf (accessed 13 July 2020).

10. CBD. 2014. Global Biodiversity Outlook 4. Montréal. https://www.cbd.int/gbo/gbo4/publication/gbo4-en.pdf (accessed 13 July 2020).

11. CBD Secretariat 2007. Submission from the Secretariat of the Convention on Biological Diversity on the Issue of Reducing Emissions from Deforestation in Developing Countries. https://unfccc.int/sites/default/files/cbd_20070227134350.pdf

12. Chandra A, Idrisova A. 2011. Convention on Biological Diversity: a review of national challenges and opportunities for implementation. Biodiversity and Conservation 20: 3295–3316. https://doi.org/10.1007/s10531-011-0141-x

13. Coad L, et al. 2013. Progress towards the CBD protected area management effectiveness targets. The International Journal for Protected Areas and Conservation 19: 13–24. https://doi.org/10.2305/IUCN.CH.2013.PARKS-19-1.LC.en

14. Di Falco S, Perrings C. 2003. Crop genetic diversity, productivity and stability of agroecosystems. A theoretical and empirical investigation. Scottish Journal of Political Economy 50: 207–216.

15. Dufresnes C, et al. 2019. Two decades of non-invasive genetic monitoring of the grey wolves recolonizing the Alps support very limited dog introgression. Scientific Reports 9: 148. https://doi.org/10.1038/s41598-018-37331-x.

16. Esquinas-Alcázar J. 2005. Protecting crop genetic diversity for food security: political, ethical and technical challenges. Nature Reviews Genetics 6: 946–953. https://doi.org/10.1038/nrg1729

17. Galli A, Wackernagel M, Iha K, Lazarus E. 2014. Ecological footprint: implications for biodiversity. Biological Conservation 173: 121–132. https://doi.org/10.1016/j.biocon.2013.10.019

18. Hoban S, et al. 2014. Comparative evaluation of potential indicators and temporal sampling protocols for monitoring genetic erosion. Evolutionary Applications 7: 984–998. https://doi.org/10.1111/eva.12197

19. Hoban S, et al. 2020. Genetic diversity targets and indicators in the CBD post-2020 global biodiversity framework must be improved. Biological Conservation 248: 108654. https://doi.org/10.1016/j.biocon.2020.108654

20. Hoffmann I. 2010. Climate change and the characterization, breeding and conservation of animal genetic resources. Animal Genetics 41: 32–46. https://doi.org/10.1111/j.1365-2052.2010.02043.x

21. Holderegger R, et al. 2019. Conservation genetics: Linking science and practice. Conservation Biology 28: 3848–3855. https://doi.org/10.1111/mec.15202

22. Hollingsworth PM et al. 2020. Scotland’s biodiversity progress to 2020 Aichi targets: conserving genetic diversity–development of a national approach for addressing Aichi Biodiversity Target 13 that includes wild species. Scottish Natural Heritage, Inverness, Scotland. https://www.nature.scot/scotlands-biodiversity-progress-2020-aichi-targets-conserving-genetic-diversity-development-national (accessed 14 August 2020)

23. Houston RD, et al. 2020. Harnessing genomics to fast-track genetic improvement in aquaculture. Nature Review Genetics 21: 389–409. https://doi.org/10.1038/s41576-020-0227-y

24. Hughes AR, Inouye BD, Johnson MTJ, Underwood N, Vellend M. 2008. Ecological consequences of genetic diversity. Ecology Letters 11: 609–623. https://doi.org/10.1111/j.1461-0248.2008.01179.x

25. IPBES, 2019. Global assessment report on biodiversity and ecosystem services of the Intergovernmental Science-Policy Platform on biodiversity and ecosystem services. Brondizio ES, Settele J, Díaz S, Ngo HT, editors. IPBES Secretariat, Bonn, Germany. https://ipbes.net/global-assessment

26. Khoury CK, et al. 2014. Increasing homogeneity in global food supplies and the implications for food security. Proceedings of the National Academy of Sciences 111: 4001–4006. https://doi.org/10.1073/pnas.1313490111

27. Khoury CK, et al. 2019. Comprehensiveness of conservation of useful wild plants: an operational indicator for biodiversity and sustainable development targets. Ecological Indicators 98: 420–429. https://doi.org/10.1016/j.ecolind.2018.11.016

28. Laikre L. 2010. Genetic diversity is overlooked in international conservation policy implementation. Conservation Genetics 11: 349–354. https://doi.org/10.1007/s10592-009-0037-4

29. Laikre L, et al. 2020. Post-2020 goals overlook genetic diversity. Science 367: 1083–1085. https://doi.org/10.1126/science.abb2748

30. Leigh DM, Hendry AP, Vázques-Domínguez E, Friesen VL. 2019. Estimated six per cent loss of genetic variation in wild populations since the industrial revolution. Evolutionary Applications 12: 1505–1512. https://doi.org/10.1111/eva.12810

31. Lotze HK, et al. 2011. Recovery of marine animal populations and ecosystems. Trends in Ecology and Evolution 26: 595–605. https://doi.org/10.1016/j.tree.2011.07.008

32. Lundmark C, et al. 2019. Monitoring the effects of knowledge communication on conservation managers’ perception of genetic biodiversity Marine Policy 99: 223–229. https://doi.org/10.1016/j.marpol.2018.10.023

33. Mathieu-Bégné E, Loot G, Chevalier M, Paz-Vinas I, Blanchet S. 2019. Demographic and genetic collapses in spatially structured populations: insights from a long-term survey in wild fish metapopulations. Oikos 128: 196–207. https://doi.org/10.1111/oik.05511

34. OECD. 2019. The Post-2020 Biodiversity Framework: targets, indicators and measurability implications at global and national level. Interim Report, November 2019. http://www.oecd.org/environment/resources/biodiversity/report-the-post-2020-biodiversity-framework-targets-indicators-and-measurability-implications-at-global-and-national-level.pdf (accessed 13 July 2020).

35. Peréz-Espona S, ConGRESS Consortium. 2017. Conservation genetics in the European Union - biases, gaps and future directions. Biological Conservation 209: 130–136. https://doi.org/10.1016/j.biocon.2017.01.020

36. Pierson JC, et al. 2016. Consideration of genetic factors in threatened species recovery plans on three continents. Frontiers in Ecology and the Environment 14: 433–440. https://doi.org/10.1002/fee.1323.

37. Potter KM, et al. 2017. Banking on the future: progress, challenges and opportunities for the genetic conservation of forest trees. New Forests 48: 153–180 https://doi.org/10.1007/s11056-017-9582-8

38. Raffard A, Santoul F, Cucherousset J, Blanchet S. 2019. The community and ecosystem consequences of intraspecific diversity: a meta-analysis. Biological Reviews 94: 648–661. https://doi.org/10.1111/brv.12472

39. Reusch TBH, Ehlers A, Hammerli A, Worm B. 2005. Ecosystem recovery after climatic extremes enhanced by genotypic diversity. Proceedings of the National Academy of Sciences 102: 2826–2831. https://doi.org/10.1073/pnas.0500008102

40. Santamaría L, Mendez PF. 2012. Evolution in biodiversity policy - current gaps and future needs. Evolutionary Applications 5: 202–218. https://doi.org/10.1111/j.1752-4571.2011.00229.x

41. Sgrò CM, Lowe AJ, Hoffmann AA. 2011. Building evolutionary resilience for conserving biodiversity under climate change. Evolutionary Applications 4: 326–337. https://doi.org/10.1111/j.1752-4571.2010.00157.x

42. Shafer ABA, et al. 2015. Genomics and the challenging translation into conservation practice. Trends in Ecology and Evolution 30: 78–87. https://doi.org/10.1016/j.tree.2014.11.009

43. Sjöqvist CO, Kremp A. 2016. Genetic diversity affects ecological performance and stress response of marine diatom populations. The ISME Journal 10: 2755–2766. https://doi.org/10.1038/ismej.2016.44.

44. Taft HR, et al. 2020. Research–management partnerships: An opportunity to integrate genetics in conservation actions. Conservation Science and Practice. https://doi.org/10.1111/csp2.218

45. Taylor HR, Dussex N, Van Heezik Y. 2017. Bridging the conservation genetics gap by identifying barriers to implementation for conservation practitioners. Global Ecology and Conservation 10: 231–242. https://doi.org/10.1016/j.gecco.2017.04.001

46. Vernesi C, et al. 2008. Where’s the conservation in conservation genetics? Conservation Biology 22: 802–804. https://doi.org/10.1111/j.1523-1739.2008.00911.x

47. Walpole M, et al. 2009. Tracking Progress Toward the 2010 biodiversity target and Beyond. Science 325: 1503–1504. https://doi.org/10.1126/science.1175466

48. Wernberg T, et al. 2018. Genetic diversity and kelp forest vulnerability to climatic stress. Scientific Reports 8: 1851. https://doi.org/10.1038/s41598-018-20009-9

