## Supplemental Information for "An analysis of genetic diversity actions, indicators and targets in 114 National Reports to the Convention on Biological Diversity"

**CBD Instructions**

Below is a table matching the instructions/ guidelines that the CBD gave countries to the questions used in our questionnaire.

The CBD instructions to countries for the 5th report are in CBD document UNEP/CBD/COP/DEC/X/10 ([link here](https://www.cbd.int/doc/decisions/cop-10/cop-10-dec-10-en.pdf)) released October 2010.

The CBD instructions to countries for the 6th report are in CBD document ([link here](https://www.cbd.int/doc/decisions/cop-13/cop-13-dec-27-en.pdf)) released in December 2016.

| **Topic** | **NR** | **CBD Instructions wording** |  | **Our question** |
| --- | --- | --- | --- | --- |
| Executive Summary | 5 | *For the purposes of communicating to stakeholders..., Parties should prepare an executive summary of the fifth national report that provides the main messages and key findings of the report… serve as a useful 'stand-alone' tool to communicate, educate and raise awareness of biodiversity among the general public, relevant decision-makers and other key stakeholder groups* |  | Q1. Is genetic diversity (e.g. genetic threats, genetics-related policy or gene conservation actions- essentially any genetic content) mentioned in the Executive Summary? |
|  | 6 | No match |  | Q10. Is genetic diversity (e.g. genetic threats, genetics-related policy or gene conservation actions- essentially any genetic content) mentioned in the Executive Summary? |
| Utility/ importance of genetic diversity | 5 | *Q1. Why is biodiversity important for your country… [including] contributions of biodiversity and related ecosystem services to human well-being and socio-economic development… AND Q4: What are the impacts of the changes in biodiversity for ecosystem services and the socio-economic and cultural implications of these impact* |  | Q2. Is the importance or utility of genetic diversity noted, and if yes, how? |
|  | 6 | *[Under the updated biodiversity country profile]...benefits from biodiversity and ecosystem services and functions* |  | Q13: Is the importance or utility of genetic diversity noted, and if yes, how? |
| Status and trends | 5 | *Q2: What major changes have taken place in the status and trends of biodiversity in your country?Focus on changes that have occurred, or that have become known, since the fourth or last national report was prepared… Where possible, show changes in biodiversity or other trends over time and use quantitative indicators* |  | Q3. Is the STATUS of genetic diversity reported on? Is the TREND of genetic diversity reported on? |
|  | 6 | *[Under the updated biodiversity country profile] Status and trends of biodiversity* |  | Q6: Is the STATUS of genetic diversity reported on? Is the TREND of genetic diversity reported on? |
| Threats/ pressures | 5 | *Q3: What are the main threats to biodiversity? (Or, what are the main causes of the negative changes described in the answer to question two?) … the main direct drivers of biodiversity loss (pressures) and the main indirect drivers (underlying causes)* |  | Q4. What genetic threats/ pressures/ drivers are reported- according to the report authors (potential or actual, measured or not)? |
|  | 6 | *[Under the updated biodiversity country profile] Main pressures on and drivers of change to biodiversity (direct and indirect):* |  | Q11: What genetic threats/ pressures/ drivers are reported- according to the report authors (potential or actual, measured or not)? |
| List of Targets | 5 | *Q5: What are the biodiversity targets set by your country?* |  | Q5. Is there a National level Target FOCUSED ON 'conserving/ maintaining genetic diversity,' 'genetic erosion,' 'genetic resources,' 'genetic loss,' etc. e.g. a national target similar to Aichi Target 13?  Q6. Is there a different national level target that REFERS to 'genetic diversity,' 'genetic erosion,' 'genetic resources,' 'genetic loss,' or other genetics-related terms or genetic data? |
|  | 6 | *Section I. Information on the targets being pursued at the national level* |  | Q1: Is there a National level Target FOCUSED ON 'conserving/ maintaining genetic diversity,' 'genetic erosion,' 'genetic resources,' 'genetic loss,' etc. e.g. a national target similar to Aichi Target 13?  Q2: Is there a different national level target that REFERS to 'genetic diversity,' 'genetic erosion,' 'genetic resources,' 'genetic loss,' or other genetics related terms or genetic data? |
| Actions taken | 5 | *Q7: What actions has your country taken to implement the Convention since the fourth report and what have been the outcomes of these actions* |  | Q7. Are there Actions planned regarding genetic diversity or Actions taken regarding genetic diversity, in or by this country? |
|  | 6 | *Section VII. Overall actions taken to contribute to the implementation of the Strategic Plan for Biodiversity 2011-2020* |  | Q3: Are there Actions planned regarding genetic diversity or Actions taken regarding genetic diversity, in or by this country? |
| Progress towards Targets | 5 | *Q10: What progress has been made by your country towards the implementation of the Strategic Plan for Biodiversity 2011-2020 and its Aichi Biodiversity Targets?* |  | Q8. What is the degree of progress reported regarding Aichi Target 13, or if Aichi progress is not reported, then progress on the National Target corresponding to the Aichi target 13, if one can be identified? |
|  | 6 | *Section III. please assess the level of progress made towards each of your country’s national targets or similar commitments.* |  | Q5: What is the degree of progress regarding outcomes of the National Target corresponding to Target 13, or if National Target progress is not reported, then progress on Aichi target 13? |
| Support/ relation to MDG/ SDGs | 5 | *Q11: What has been the contribution of actions to implement the Convention towards the achievement of the relevant 2015 targets of the Millennium Development Goals in your count* |  | Q9. Is genetic diversity (e.g. genetic data, policy, threats, actions) mentioned under global sustainable development initiatives (MDGs/ SDGs)? |
|  | 6 | *Section IV. please describe how and to what extent these contributions support the implementation of the 2030 Agenda for Sustainable Development and the Sustainable Development Goals:* |  | Q7: Is genetic diversity (e.g. genetic data, policy, threats, actions) mentioned under global sustainable development initiatives (MDGs, SDGs)? |
| Lessons Learned & Obstacles/ needs | 5 | *Q12: What lessons have been learned from the implementation of the Convention in your country?* |  | Q10. What lessons were learned about genetic diversity if any?  Q11. Are obstacles or needs (e.g. technological, scientific, etc.) listed for the genetic diversity target (e.g. Aichi Target 13 or National Target on genetic diversity)? |
|  | 6 | *Section II. Please describe what obstacles have been encountered and any scientific and technical needs for addressing these* |  | Q4: Are obstacles or needs (technological, scientific, etc.) listed for the genetic diversity target (e.g. Aichi Target 13 or National Target on genetic diversity)? |
| GMOs | 5 | *None* |  | Q12. What is the general tone regarding Genetically Modified Organisms, or GMOs? |
|  | 6 | *None* |  | Q12. What is the general tone regarding Genetically Modified Organisms, or GMOs? |
| Relation to indigenous people and local communities | 5 | *None* |  | Q13. Is genetic diversity mentioned with respect to indigenous peoples and local communities, including contributions towards their conservation, use of, or their value of genetic diversity? |
|  | 6 | *Section VI. Additional information on the contribution of indigenous peoples and local communities to the achievement of the Aichi Biodiversity Targets* |  | Q9: Is genetic diversity mentioned with respect to indigenous peoples and local communities, including contributions towards their conservation, use of, or their value of genetic diversity? |
| GSPC | 5 | *None* |  | None |
|  | 6 | *Section V. describe your country’s contribution towards the achievement of the targets of the Global Strategy for Plant Conservation… [including] the major measures taken by your country for the implementation of the Global Strategy for Plant Conservation.* |  | Q8: What is the progress on genetic diversity related Targets of the Global Strategy for Plant Conservation? |
