## Supplemental Information for "An analysis of genetic diversity actions, indicators and targets in 114 National Reports to the Convention on Biological Diversity"

**This document contains Supplemental Methods, Results and Discussion for the article “Reporting to the Convention on Biological Diversity: How are countries assessing and protecting their genetic diversity?”**

Table of contents

-Supplemental Methods

-Supplemental Results, summary statistics

-Significance Tests comparing 5th and 6th Reports

-Significance Tests for Socio-economic status

-Complete tabulation of results for each test

-Supplemental Expanded Discussion

**Supplemental Methods**

**Questionnaire development.**

We composed the questions and detailed instructions on how to answer the questions via several phases.

Phase 1, scoping and first draft: First, we read numerous National Reports to understand their format and content. We developed a draft set of questions that required reviewers to input short answers via a spreadsheet format. Our team reviewed a set of reports to determine if this format would suffice; this scoping exercise took place from late January to March 2019. We considered two types of questions: general questions about genetic diversity content and questions about the “types” of species that were mentioned in the Reports. A total of 40 Reports were reviewed, focused more on the 5th Report as more of these were available at the time (most countries submit reports late, up to several years late), but also including five paired (from the same country) 5th and 6th National Reports. Through several conference calls we discussed the outcome- a large variance in how reviewers interpreted questions and the level of detail used in filling in spreadsheets, such as whether short answers, extensive quoting, or long answers were used. Therefore, we determined that a standard set of question options (e.g. check boxes) for reviewers to choose from, rather than short answers, would be needed.

Phase 2, a new draft: Next, a core group of authors composed standard answer choices for each question via a series of four group discussion calls in April and May 2019, after which a second set of authors commented on and suggested revisions to the questions, answers and instructions that include clear definitions of terminology. Several additional group conversations took place to determine further instructions for reviewers. A separate small group volunteered to revise the question spreadsheets. All authors then provided input on this near-final set of questions.

Phase 3, further testing and revising: After finalizing the questions and answers, we reviewed 15 reports with two reviewers per report to determine consistency among reviewers’ answers. This was started in August 2019. We focused on a small number of countries, making sure that these countries were reviewed by two separate people. At the end of August we qualitatively and quantitatively assessed agreement between the primary and secondary reviewers. We determined that we could make some further improvements in instructions to the reviewers and small changes in the question wording. This took place in late August/ early September 2019. Each pair of reviewers had a one-on-one conversation to discuss discrepancies between their answers and their logic, and arrive at a final set of answers. They made recommendations to improve clarity of the questionnaire, primarily by adding additional instructions and some examples. Following this comparison, we again further revised the questions and instructions to reviewers on the basis of common differences in interpretation. The final set of questions is in Supplemental Document B, which shows the CBD language that helped develop each of our questions. The full instructions to reviewers can be found In Supplemental Docs C through E.

Phase 4, data collection: Data collection began again in September 2019. All reviewers were instructed to read the new instructions and questions in detail. For those reports which were previously reviewed by one reviewer, the reviewer was to update their answers based on new instructions. For those reports that were reviewed in duplicate, the two reviewers had a face to face call to come to an agreement and then update the answers,on the basis of the agreement and on the new instructions. Most reports, however, were being reviewed for the first time. All questionnaires were completed by January 2020.

**Agreement among reviewers.** The rate of agreement between reviewers was assessed by counting the number of individual responses that were the same against the total number of responses. “Same response” is counted when both reviewers check the same box or when both reviewers do not check that box. Disagreement is when one reviewer checks a box and the other does not. For the questions with the subcategory Qualitative and Quantitative, because we do not focus on Quantitative vs. Qualitative responses, we considered the reviews in agreement if either of these was checked by both reviewers, e.g. if one checked Qualitative and one Quantitative we consider this still agreement. The following questions were considered for the calculation: Status/ Indicator used (3 parts), Actions (2 parts), Progress to Target, SDG mentioned, GMO mention, indigenous uses mentioned, Uses, Threats, mention in Executive Summary (only for 5th report), and Targets.

**Reports we analyzed.** The countries whose National Reports we could review were determined as all countries that had submitted their Sixth National Reports as of 1 July 2019, and also had a Fifth National Report available, which was 63 countries. One country was removed as we had no fluent Russian speakers, another country was removed as its report was too short (2 pages) and a further 4 countries were removed due to time constraints. As such, the fifth and sixth reports from 57 countries were included in this study. A list of all countries included and a map is provided below, showing the global scope.

Supplemental Table S1: All countries included in the study which has both 5th and 6th National Reports analyzed.

| Andorra | Dominican Republic | Jordan | Norway | Sweden |
| --- | --- | --- | --- | --- |
| Antigua & Barbuda | Ecuador | Kazakhstan | Panama | Switzerland |
| Belgium | Estonia | Korea | Peru | Thailand |
| Bhutan | Ethiopia | Kyrgyzstan | Philippines | Togo |
| Botswana | European Union | Lebanon | Poland | Tunisia |
| Burkina Faso | Finland | Luxembourg | Moldova | United Kingdom |
| Cameroon | France | Mali | Saint Kitts and Nevis | Tanzania |
| Canada | Gambia | Mexico | Slovakia | Uruguay |
| Chile | India | Morocco | South Africa | Zambia |
| China | Italy | Myanmar | Spain |  |
| Costa Rica | Cote d'Ivoire | Nepal | Saint Vincent and the Grenadines |  |
| Czech Republic | Japan | Niger | Sudan |  |

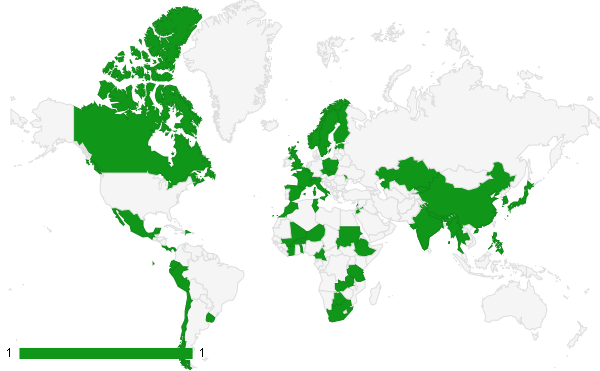

Supplemental Figure S1: Map of countries included in this study.

**Additional analyses of responses categorized as “other”**. A relatively small number of respondents chose “other” for several questions. “Other” responses allowed a text box description. We performed a post hoc categorization of these “other” responses into the existing categories, placing the response in a category by means of consensus among three authors, when possible, though some “other” responses were truly other and could not be categorized. This was performed for the questions on actions, threats, and indicators. When “other”responses that were recategorized are analyzed in the results, the findings do not change substantially, though small differences are observed, as follows:

For **actions**, Fisher’s exact test results do not change (i.e. not significant), while the t-test shows stronger significance (0.056 instead of 0.11), for a higher number of actions in the 6th National Report. The plots without (top plots) and with (bottom plot) these new responses are shown below, demonstrating high similarity in trends.

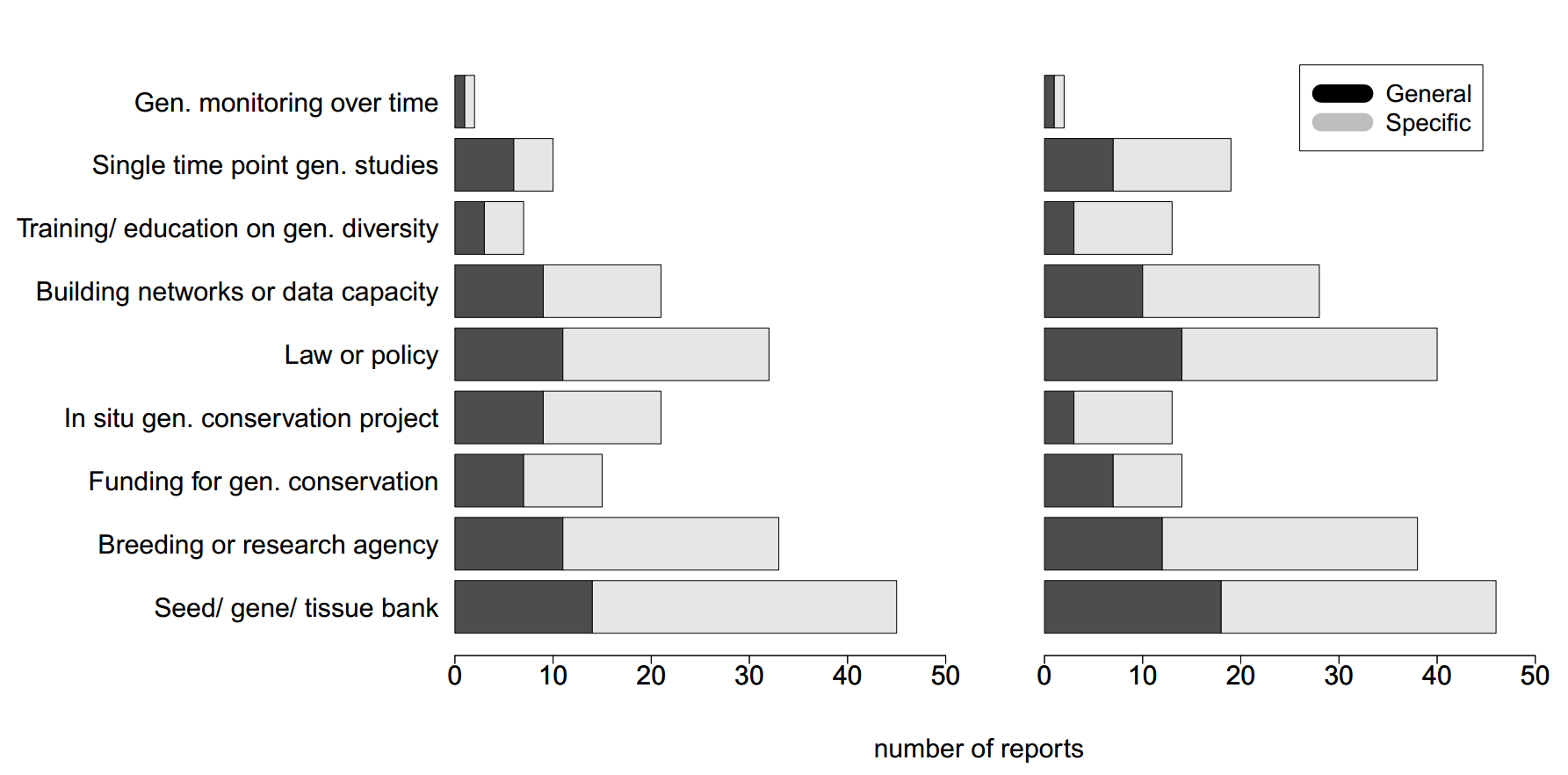

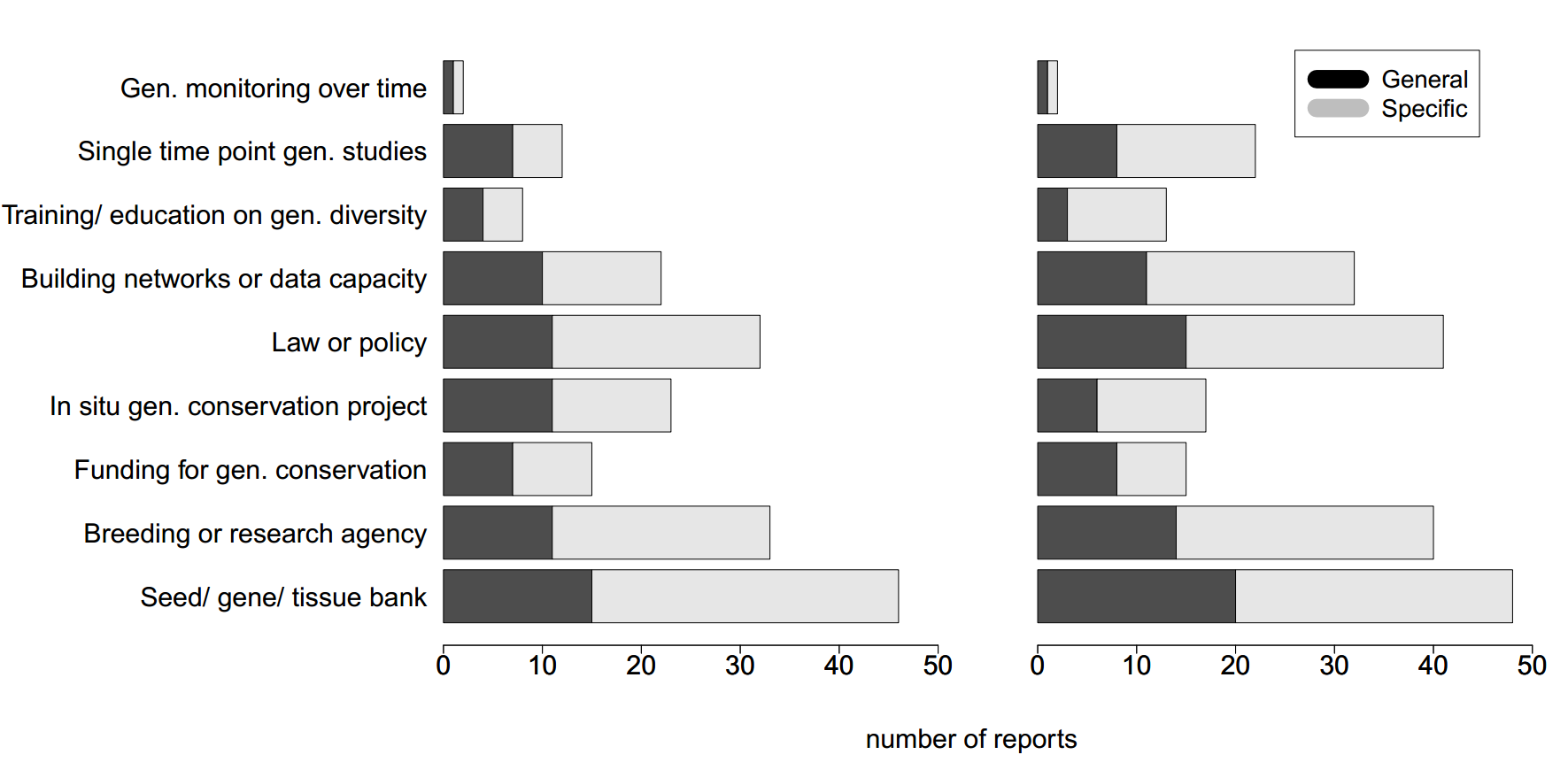

For **threats**, Fisher’s exact test results do not change i.e. not significant), and the T-test essentially does not change (0.04 instead of 0.02), for a higher number of threats in the 5th National Report. The plots without (top plots) and with (bottom plot) these new responses are shown below, demonstrating high similarity in trends.

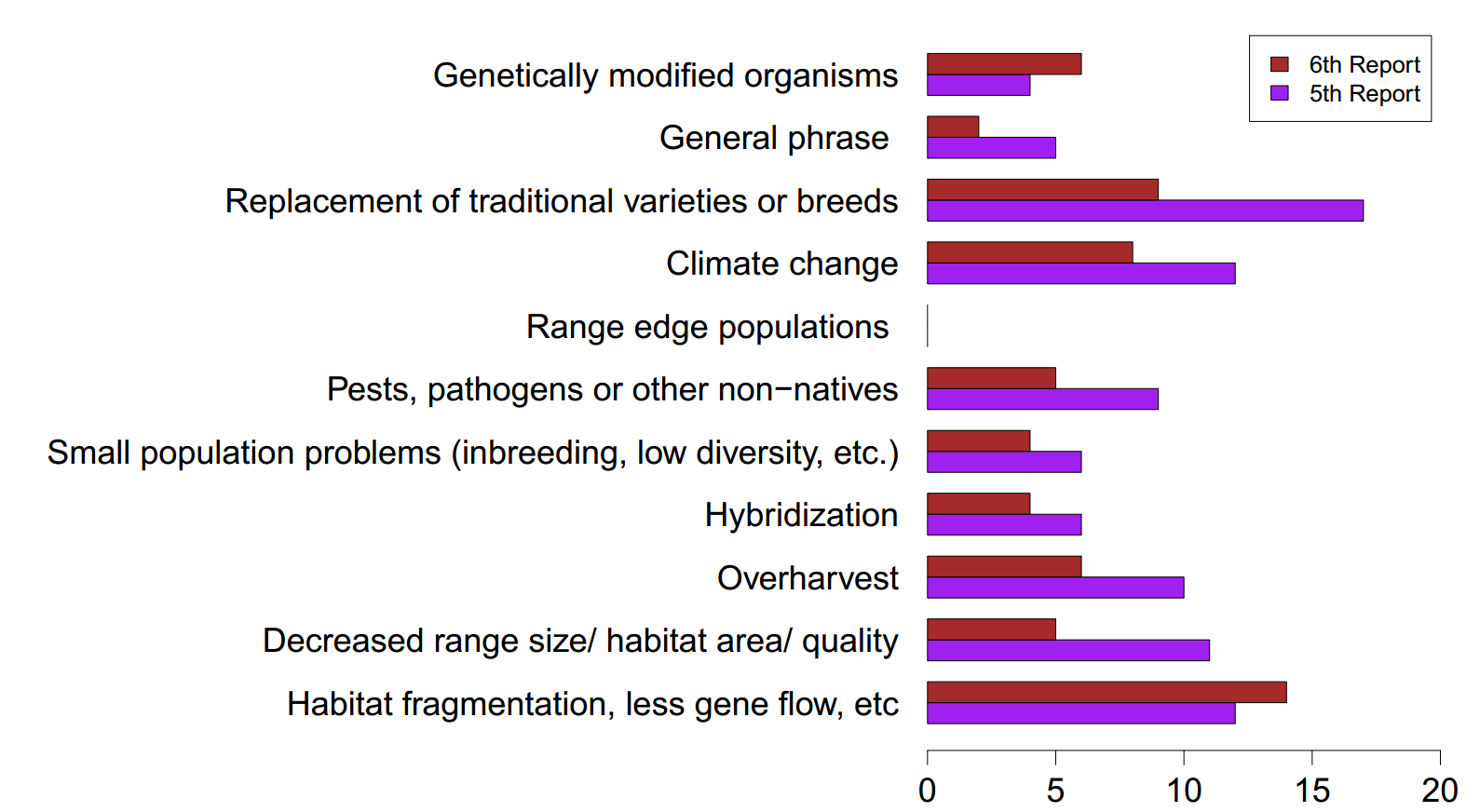

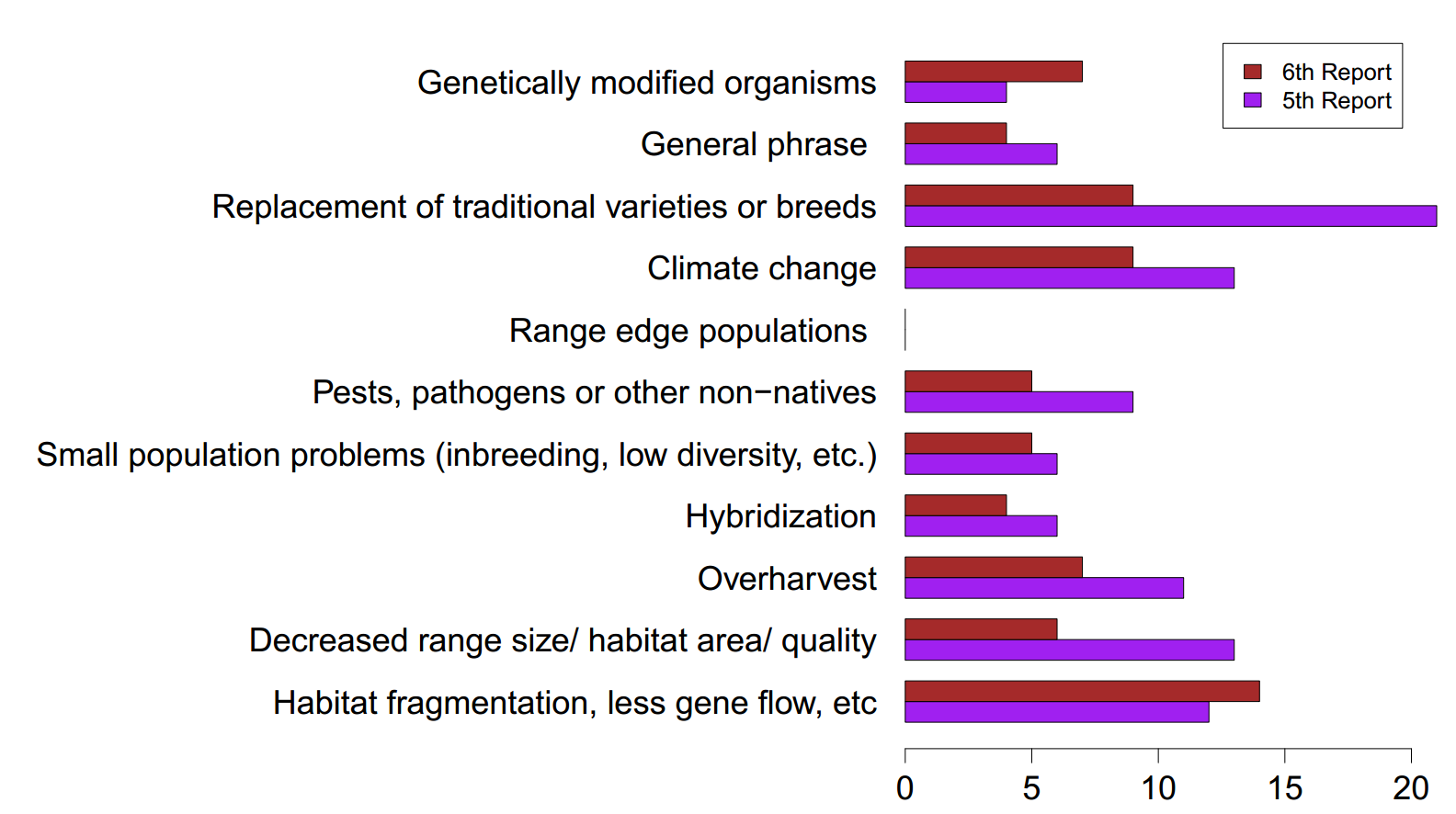

For **indicators**, the Fisher’s exact test results do not change (i.e. not significant), nor does the Wilcoxon test (p value remains the same at 0.106). The plots without (top plots) and with (bottom plot) these new responses are shown below, demonstrating high similarity in trends.

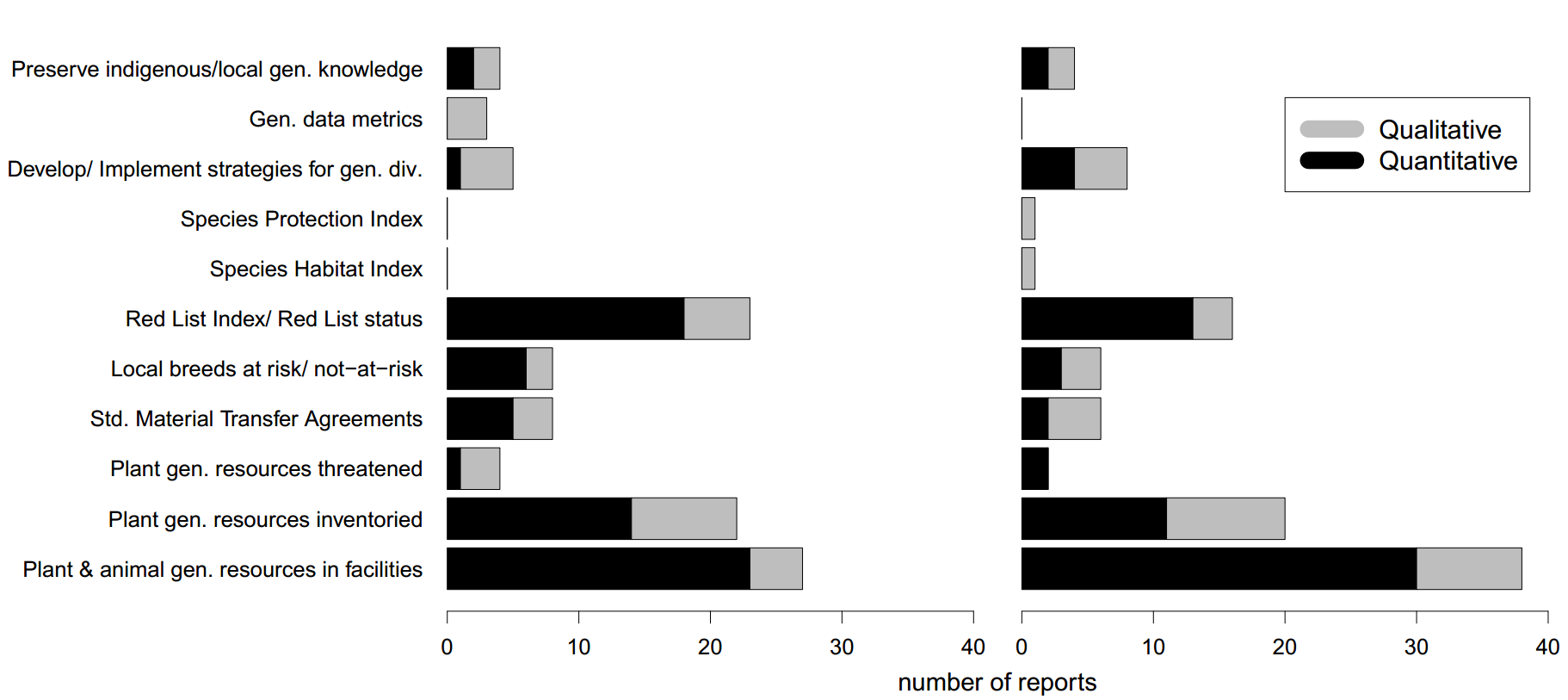

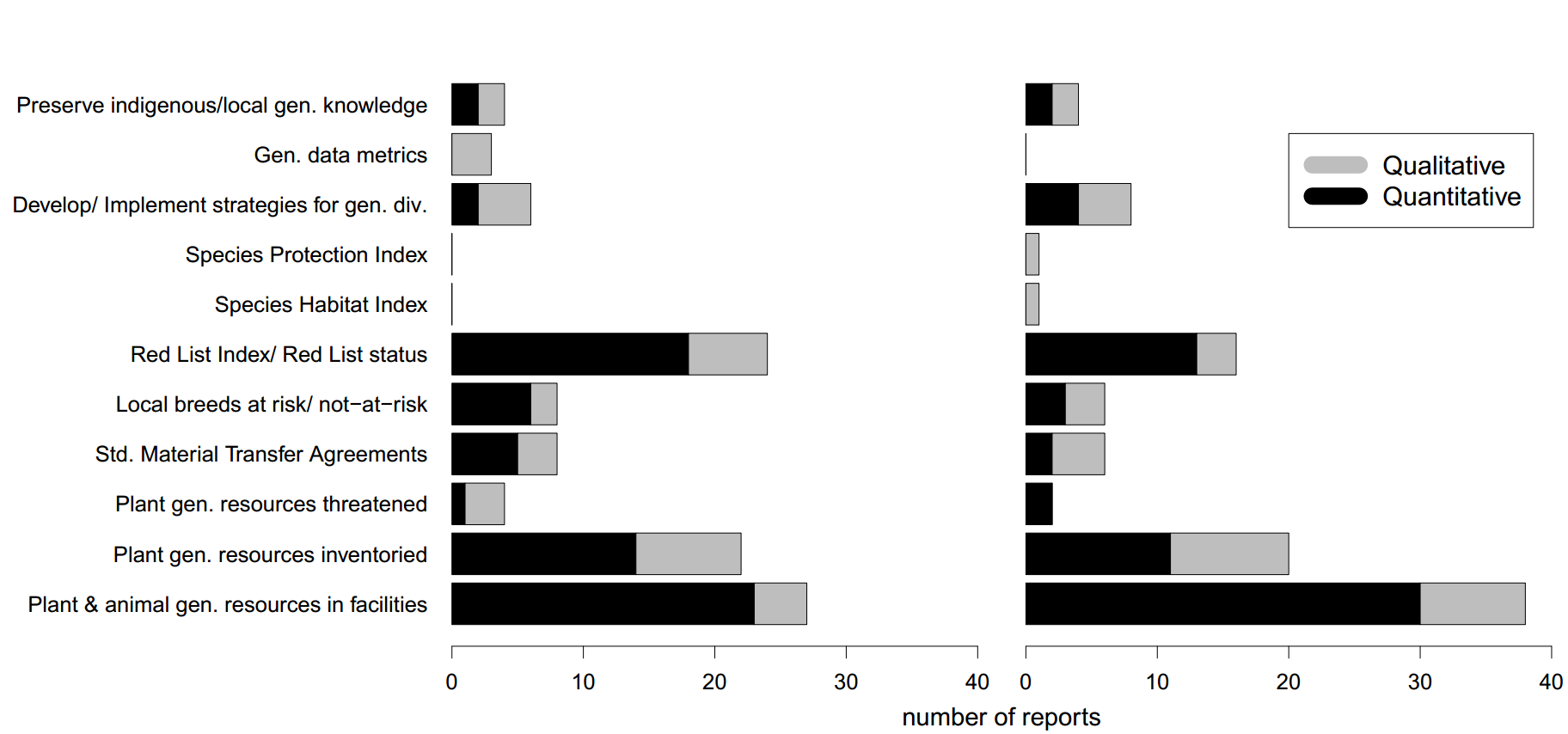

**Supplemental Results- basic statistics**

**General summary- length, agreement.** The median length of the 57 reports analyzed was 49,428 words (5th Report) and 52,712 words (6th Report), with the 6th National Reports being 24% longer (paired t test, p=0.01). The 57 reports analyzed were slightly longer than all published National Reports (5th Report median 42,451; 6th Report 51,341). Length was correlated between the Reports (countries with long 5th reports tended to have long 6th Reports: rho=0.562, p<0.001). Medium income countries have the longest reports. Fifteen reports were reviewed separately by two reviewers. The lowest agreement regards the Uses/ Values of genetic diversity question, thus we emphasize that higher caution should be used for interpreting this question and we do not perform formal analysis of this question. Excluding this category, agreement was 89% for the 5th report and 85% for the 6th report.

**Supplemental Results- Compare 5th to 6th Reports**

**Significance of comparisons 5th vs. 6th**. As explained in the Methods, we performed two types of tests- (1) we tested whether the frequency distribution of responses differed between reports, and (2) we tested whether the total count of responses differed between reports

(1) To compare the frequency distribution of responses in each category of answers (for example, the categories of types of Actions) for each question in the 5^th^ and 6^th^ report we used Fisher’s exact tests. These contingency tests for comparing 5th and 6th Reports were largely not significant (e.g. no temporal change in proportion of responses in each answer category), all p values are given below
(2) To compare the mean number of responses recorded (for example the number of Actions) between the 5th and 6th report we used paired t tests when data were normally distributed and Wilcoxon signed rank tests when not. Below we present p values for every comparison made between 5th and 6th National Reports, with a summary table at the end.

**Values:** The proportions of each category of value did not change between the 5th and 6th Report (Fisher’s exact test, p=0.72). There were 34% more mentions of values of genetic diversity in the 5^th^ National Report compared to the 6th report (p=0.04).

**Target 13 or equivalent:** There was no significant difference between the 5th and 6th Reports in whether the country had a genetic diversity Target (Fisher’s exact test p=0.35), though there was a slight increase (40 for the 5th Report and 45 for the 6th). There was also no significant difference between reports in proportion that could refer to non agricultural and non economically important species (Fisher’s exact test, p=0.51).

**Progress:** There is no statistically significant difference in the proportion of countries in each progress category between the 5th and 6th report (Fisher’s exact test, p=0.469). Note that progress was much more frequently reported (or clearly identifiable) in the 6^th^ Report, probably because it was only in the 6th report that the CBD listed these clear categories.

**Other Targets and genetic diversity:** Countries referenced genetics in relation to 13 different (non genetic) Aichi Targets in the Fifth report, but 19 of them in the Sixth report; a significant increase (Fisher’s exact test p=0.044) of almost 50%. The number of countries referencing genetic diversity in non Target 13 and 16 Targets also increased. In the Fifth report iteration, only 11 out of 57 countries (19.3 percent) included references to genetics associated with Aichi targets other than 13 and 16, but 28 out of 57 (49.1 percent) of the surveyed 6th Reports mentioned genetics in targets other than 13 and 16, more than 2.5 times as many, a significant increase (p=0.001).

**Indicators of status and trends:** The proportion of each category of indicator did not have significant differences between the 5th and 6th report (chi square test, p=0.12). There were similar numbers of mentions of indicators of status of genetic diversity in the 5th and 6th report (paired t-test: p=0.337). Trends were noted 41% more in the 5th Report than the 6th (Wilcoxon test: p=0.106).

**Actions:** There is no significant difference between the 5th and 6th report among categories of actions (chi square test, p=0.43), nor is there a significant difference between “specific” and “general” actions (see Table 1) within a reporting period (chi square test, 5th Report, p=0.15; 6th Report, p=0.46). The most and least commonly reported actions did not significantly change over time, or in terms of “specific vs. “general” reporting. There were slightly more actions (17% more, per category) identified in the 6^th^ than the 5^th^ Report (paired t-test, marginally not significant p=0.111). The rank order of the status indicators does not differ between reports, and the percent differences are modest (see Table below).

**Threats:** The percentages in each threat category did not differ between the 5th and 6th reports (Fisher’s exact test, p=0.86). There were about 46% more threats mentioned in the 5^th^ National Report (i.e. there was a decrease in these mentions in 6th Report, paired t-test p=0.02

**Species Types:** In the categorization of the species mentioned, there were no significant differences between categories of threat, action, change and status within the 5th and within the 6^th^ Reports (Chi square test p = 0.90 and 0.96, respectively), or between the reporting periods (Fisher’s exact test p=0.12). There were slightly (17%) more species mentions in the 5th report (Wilcoxon test, p =0.256).

**Summary of differences between the 5th and 6th reports**. There is more mention of genetic diversity with respect to genetic diversity under other National Targets for the 6th Report, possibly due to increasing awareness of genetic diversity and increased consideration of Target 13. However the 5th Report had more mentions of values, trends, threats, and mentions of specific species, possibly because it was a more narrative, less structured report.

Supplemental Table S2: Significance values (P) for all comparisons between the 5th and 6th National Reports

| **Question/ category** | **Summary** | **Which had more?** | **P** |
| --- | --- | --- | --- |
| Executive summary | Only asked for in the 6th Report | N/A |  |
| Values/ Uses/ Importance | --Slightly more values mentions in 5th Report  --No difference in proportions of each category | 5th, 34%  N/A | 0.04  n.s. |
| Genetic diversity National Target | --Slightly higher % of countries in 6th Report with genetic diversity National Target  --Higher % of National Targets in 6th Report that include non agricultural species - | 6th, slight  6th, slight | n.s.  n.s |
| Progress on Target | --No significant difference among categories of progress | No diff | n.s. |
| Genetic diversity under other Targets | --More countries mentioning genetic diversity under other Targets for 6th Report  --More other Targets mentioning genetics | 6th, 250%  6th, 50% | 0.001  0.044 |
| Status | --Similar numbers of mentions 5th vs 6th  --Slight changes in proportions for each category of Status | No diff  N/A | n.s  n.s. |
| Trends | --More reports of trends in 5th Report (nearly signif p=0.08) | 5th, 41% | 0.106 |
| Actions | --More actions in the 6th Report (nearly signif)  --No difference in proportions of each category | 6th, 17%  N/A | n.s.  n.s. |
| Threats | --More threats mentioned in 5th Report (p=0.02)  --No difference in proportions of each category | 5th, 46%  N/A | 0.02  n.s. |
| Species mentions | --Slightly more mentions of species in 5th Report (p=0.10)  --No difference in proportions of each category | 5th, 17% | n.s.  n.s. |

**Supplemental Results- Socio economic status**

**Details on differences between countries’ economic status**

Low-income and middle-income countries were more likely than high-income countries to mention genetics in relation to genetic diversity targets, genetic resources targets and in targets outside of target 13 and 16, though none of these results are significant. Specifically, low-income and middle-income countries were more likely than high-development countries to mention genetics in relation to targets outside of target 13 and 16 (5th Report: low: 21.1% and middle: 26.7%, respectively, compared to high: 13.0%; 6th Report: 63.2%, 40.0% and 43.5%) though these are not significant (5th Report: p=0.34; 6th Report: p=0.37). In the 6th Report, low-income and middle-income countries were more likely to include a Target 16 equivalent (e.g. access and benefits to genetic resources) than high-income countries (73.7% of low-income, 53.3% of middle-income, and 52.2% of high-income countries did so; p=0.44). Moreover, lower and middle income countries were more likely to include a genetic diversity Target (e.g. a Target 13 equivalent) compared to high-development countries, though not significantly so (100% low-income countries, 66.7% for middle-income countries and 69.6% for high-income countries; p=0.25).

**Additional Results Tables and Figures**

**TABULATED RESULTS ON INCLUSION OF GENETIC DIVERSITY IN EXECUTIVE SUMMARY**

Supplemental Table S3: Count and proportions of responses for each question option regarding the Executive Summary

| **Genetics related text in Executive Summary** | **5^th^ NR** | **proportion** |
| --- | --- | --- |
| no executive summary or cannot answer | 10 | 0.18 |
| mention of biotechnology or access/ benefits | 17 | 0.30 |
| mentioned with other components (e.g. species) | 9 | 0.16 |
| no mention | 4 | 0.07 |
| partial reference (traits, gene banks, breeds) | 28 | 0.49 |
| direct mention genetic diversity e.g. at population genetic level | 18 | 0.32 |

*Note the Executive Summary is not found in more than 40% of 6^th^ Reports so is not included here. Also proportions add up to more than one because more than one option can apply to a given report

**TABULATED DATA ON VALUES/ USES/ IMPORTANCE OF GENETIC DIVERSITY**

Supplemental Table S4: Count and proportions of responses for each question option regarding the values or importance of genetic diversity. Data are sorted according to mean proportion across time periods

|  | **5th count data** | **5th proportion** | **6th count data** | **6th proportion** | **Mean proportion** |
| --- | --- | --- | --- | --- | --- |
| resilience to climate, drought, other env. change | 21 | 0.368 | 13 | 0.228 | 0.298 |
| increase productivity (forest, fishery, agriculture) etc. | 21 | 0.368 | 13 | 0.228 | 0.298 |
| other direct use e.g. new varieties | 15 | 0.263 | 13 | 0.228 | 0.246 |
| adaptation via natural selection in changing environments | 15 | 0.263 | 11 | 0.193 | 0.228 |
| important but no specifics | 10 | 0.175 | 11 | 0.193 | 0.184 |
| ecosystem stability, services, or human well being | 9 | 0.158 | 4 | 0.070 | 0.114 |
| resilience in a general way | 5 | 0.088 | 7 | 0.123 | 0.105 |
| reduce “inputs” for production | 7 | 0.123 | 4 | 0.070 | 0.096 |
| avoid inbreeding in populations | 4 | 0.070 | 4 | 0.070 | 0.070 |

*Note proportions add up to more than one because more than one option can apply to a given report

**TABULATED RESULTS ON GENETIC DIVERSITY NATIONAL TARGET WORDING**

Supplemental Table S5: Count and proportions of responses for each question option regarding the National Targets. Data are sorted according to mean proportion across time periods

| **Genetic diversity Target and type of species for Target if mentioned** | **5^th^ count data** | **5th proportion** | **6^th^ count data** | **6th proportion** | **mean proportion** |
| --- | --- | --- | --- | --- | --- |
| No genetic diversity National Target | 14 | 0.250 | 9 | 0.160 | 0.205 |
| Genetic target but no specification of species use | 9 | 0.160 | 0 | 0.000 | 0.080 |
| Genetic target, only agricultural/ domesticated spp | 13 | 0.230 | 12 | 0.210 | 0.220 |
| Genetic target, alludes to non agricultural e.g., socio economic spp | 11 | 0.190 | 13 | 0.230 | 0.210 |
| Genetic target, alludes to non agricultural & non socio economic spp | 10 | 0.180 | 18 | 0.320 | 0.250 |
| Unable to determine | 3 | 0.050 | 3 | 0.050 | 0.050 |

**TABULATED RESULTS ON PROGRESS ON GENETIC DIVERSITY TARGET**

Supplemental Table S6: Count and proportions of responses for each question option regarding the progress towards the Target. Data are sorted according to mean proportion across time periods

| **Status** | **5th count data** | **5th proportion*** | **6th count data** | **6th proportion*** | **mean proportion** |
| --- | --- | --- | --- | --- | --- |
| Progress = On track to exceed | 1 | 0.040 | 3 | 0.070 | 0.055 |
| Progress = On track to achieve | 7 | 0.300 | 17 | 0.380 | 0.340 |
| Progress = Some progress but insufficient | 13 | 0.570 | 20 | 0.440 | 0.505 |
| Progress = No change | 1 | 0.040 | 6 | 0.130 | 0.085 |
| Progress = Moving away from Target | 1 | 0.040 | 0 | 0.000 | 0.020 |
| Progress = Unknown | 0 | 0.000 | 0 | 0.000 | 0.000 |
| Cannot interpret or no measure given | 28 |  | 7 |  |  |
| No Aichi Target included | 4 |  | 2 |  |  |

*proportion of those that actually reported a clear measure of progress- e.g. excludes last two rows of the table

**TABULATED RESULTS ON GENETIC DIVERSITY IN OTHER NATIONAL TARGETS**

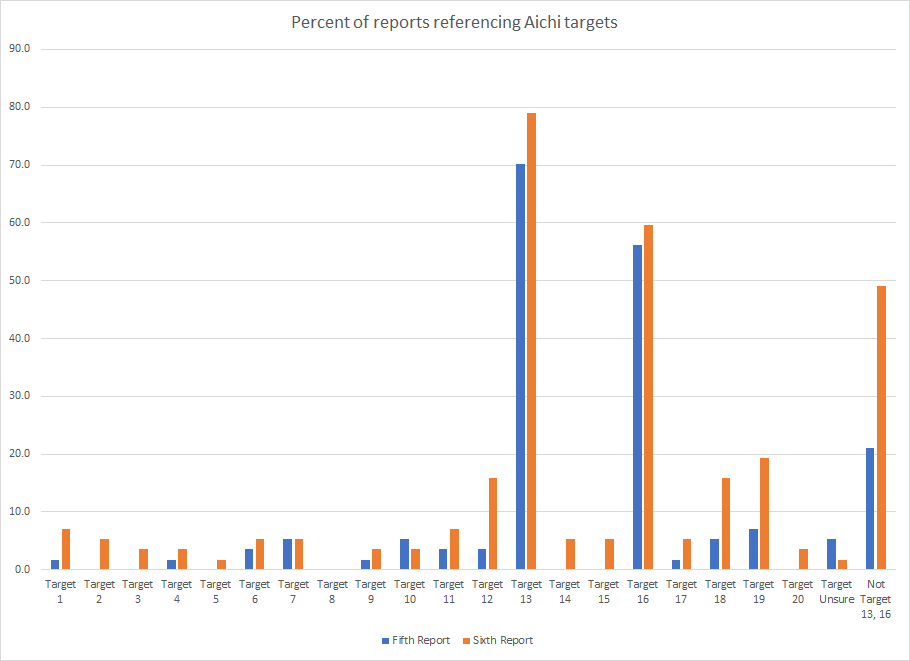

Supplemental Figure 2: Percent of nations referencing genetics in relation to each of the 20 Aichi Biodiversity Targets, for the 5th and 6th reports.

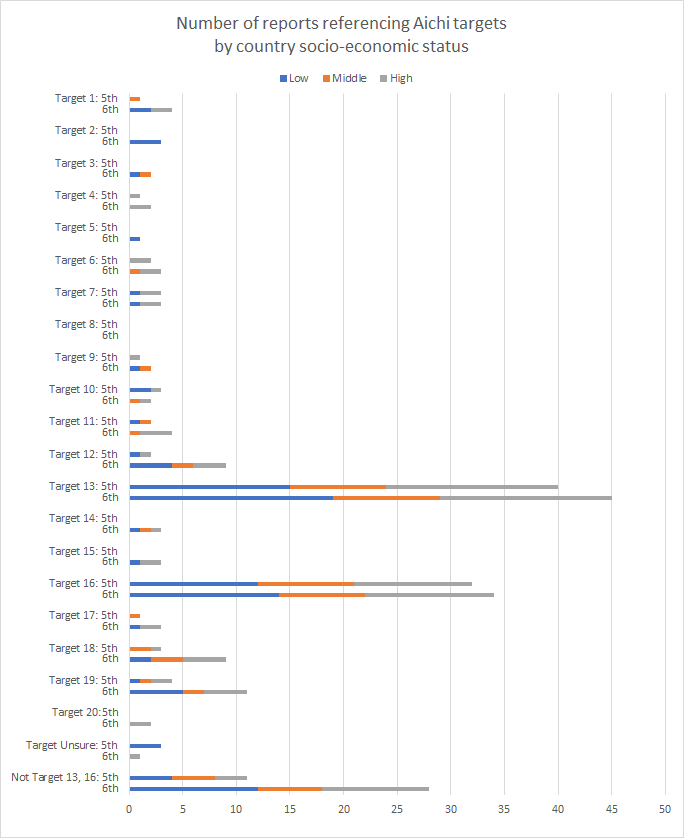

Supplemental Figure 3: Number of nations referencing genetics in relation to each of the 20 Aichi Biodiversity Targets, for the Fifth and Sixth reports, by socio-economic status

**TABULATED RESULTS ON INDICATORS OF STATUS**

Supplemental Table S7: Counts of responses for each question option regarding the indicators reported for genetic status.

|  | **Quant 5** | **Qual 5** | **Quant 6** | **Qual 6** |
| --- | --- | --- | --- | --- |
| Number of plant, animal gen resources in facilities | 23 | 4 | 30 | 8 |
| Number of plant genetic resources inventoried | 14 | 8 | 11 | 9 |
| Red List Index/ Red List status | 18 | 5 | 13 | 3 |
| Standard Material Transfer Agreements | 6 | 2 | 3 | 3 |
| Number of local breeds at risk/not-at-risk | 5 | 3 | 2 | 4 |
| Develop/ Implement strategies for... gen div | 1 | 4 | 4 | 4 |
| Preserve indigenous/ local gen div knowledge | 2 | 2 | 2 | 2 |
| Num. of plant genetic resources threatened | 1 | 3 | 2 | 0 |
| Genetic data on any species | 0 | 3 | 0 | 0 |
| Species Habitat Index | 0 | 0 | 0 | 1 |
| Species Protection Index | 0 | 0 | 0 | 1 |

**TABULATED RESULTS ON INDICATORS OF CHANGE/ TRENDS**

Supplemental Table S8: Counts of responses for each question option regarding the indicators reported for genetic trends.

Decr = decreasing, Incr = Increasing. Note that “Decreasing” means moving in an undesirable state for biodiversity conservation, e.g. the Red List Index is moving towards Critically Endangered or fewer accessions are preserved in seed banks, but also more local breeds being endangered. “Increasing” means moving in a desired state for conservation, e.g. the Red List Index is moving towards Least Concern or more accessions are preserved in seed banks, but also fewer local breeds being endangered (larger population sizes for the breeds).

|  | 5th report |  |  | 6th report |  |  |
| --- | --- | --- | --- | --- | --- | --- |
|  | Decr | Incr | No change | Decr | Incr | No change |
| Number of plant, animal gen resources in facilities | 1 | 15 | 1 | 2 | 13 | 1 |
| Number of plant genetic resources inventoried | 4 | 6 | 0 | 1 | 4 | 2 |
| Num. of plant genetic resources threatened | 1 | 0 | 0 | 1 | 0 | 0 |
| Number of local breeds at risk/not-at-risk | 3 | 1 | 0 | 1 | 0 | 0 |
| Standard Material Transfer Agreements | 0 | 4 | 1 | 0 | 1 | 0 |
| Red List Index/ Red List status | 6 | 5 | 1 | 2 | 2 | 1 |
| Species Habitat Index | 0 | 0 | 0 | 0 | 0 | 0 |
| Species Protection Index | 0 | 0 | 0 | 0 | 0 | 0 |
| Develop/ Implement strategies for... gen div | 0 | 4 | 1 | 0 | 2 | 0 |
| Genetic data on any species | 1 | 1 | 1 | 0 | 0 | 0 |
| Preserve indigenous/ local gen div knowledge | 0 | 0 | 0 | 0 | 4 | 0 |

**TABULATION OF THREATS TO GENETIC DIVERSITY**

Supplemental Table S9: Counts and proportions of responses for each question option regarding the threats reported for genetic diversity.

|  | **5th report** | **proportion** | **6th report** | **proportion** |
| --- | --- | --- | --- | --- |
| Habitat fragmentation, less gene flow, etc | 12 | 0.211 | 14 | 0.246 |
| Decreased range size/ habitat area/ quality | 11 | 0.193 | 5 | 0.088 |
| Overharvest | 10 | 0.175 | 6 | 0.105 |
| Hybridization | 6 | 0.105 | 4 | 0.070 |
| Small population problems (inbreeding, low diversity, etc.) | 6 | 0.105 | 4 | 0.070 |
| Pests, pathogens or other non-natives | 9 | 0.158 | 5 | 0.088 |
| Range edge populations | 0 | 0.000 | 0 | 0.000 |
| Climate change | 12 | 0.211 | 8 | 0.140 |
| Replacement of traditional varieties or breeds | 17 | 0.298 | 9 | 0.158 |
| General phrase | 5 | 0.088 | 2 | 0.035 |
| Genetically modified organisms | 4 | 0.070 | 6 | 0.105 |

*Note proportions add up to more than one because more than one option can apply to a given report

**TABULATION OF ACTIONS FOR GENETIC DIVERSITY**

Supplemental Table S10: Counts of responses for each question option regarding the actions reported for genetic diversity.

|  | General | Specific | General | Specific |
| --- | --- | --- | --- | --- |
| Seed/ gene/ tissue bank | 14 | 31 | 18 | 28 |
| Law or policy | 11 | 21 | 14 | 26 |
| Breeding or research agency | 11 | 22 | 12 | 26 |
| Building networks or data capacity | 9 | 12 | 10 | 18 |
| In situ gen. conservation project | 9 | 12 | 3 | 10 |
| Funding for gen. conservation | 7 | 8 | 7 | 7 |
| Single time point gen. studies | 6 | 4 | 7 | 12 |
| Training/ education on gen. diversity | 3 | 4 | 3 | 10 |
| Gen. monitoring over time | 1 | 1 | 1 | 1 |

**TABULATION OF MENTIONS OF SPECIES TYPES**

Supplemental Table S11: Counts and proportions of responses for each question option regarding the indicators reported for genetic status.

|  | 5th Report Count* | Prop | 6th Report Count* | Prop |
| --- | --- | --- | --- | --- |
| Crops | 92 | 0.261 | 72 | 0.228 |
| Farmed animals | 72 | 0.204 | 69 | 0.218 |
| Crop wild relatives | 33 | 0.093 | 43 | 0.136 |
| Species of conservation concern | 40 | 0.113 | 30 | 0.095 |
| Forestry/ logging species | 34 | 0.096 | 35 | 0.111 |
| Horticultural species | 11 | 0.031 | 21 | 0.066 |
| Wild relatives of domesticated animals | 12 | 0.034 | 9 | 0.028 |
| Culturally important species | 15 | 0.042 | 6 | 0.019 |
| Wild harvested plants | 17 | 0.048 | 17 | 0.054 |
| Species providing ecosystem services | 13 | 0.037 | 3 | 0.009 |
| Wild harvested animals | 14 | 0.040 | 11 | 0.035 |

Count refers to the sum of mentions with respect to actions, threats, status and change. Prop is a proportion of all the times that species were mentioned and thus adds up to 1.

Supplemental Expanded Discussion

**Caveats:** First, we did have to exclude several countries due to time constraints and language; however our ability to include three major global languages makes our work broader than many conservation efforts that only focus on English language. The design of our questionnaire was subjective, though it was based on the experience of experts/authors who work in situ and ex situ, in academia and beyond, and with a variety of ecosystems and wild and domesticated species. There are doubtless other categories of, for example, Actions that we did not include in our survey. Also, despite our focus on unambiguous questions, there remains subjectivity in interpreting the reports (which are themselves interpretations of other reports, data, personal communications etc).

Report writers may lack sufficient knowledge to interpret and synthesize genetic information. The national editors may have access to specialists, but may not themselves have backgrounds in biodiversity or even science, and final approval of reports is typically given by a politician or government official.

Reports may be biased in what is reported and writers typically select a few cases to highlight progress or challenges within the country (with limited published documentation to support statements).The broad scope, along with limited time and financial resources, means that it is impractical to include all threats, actions and assessments of their effectiveness. The reports are at best summaries, which represent the areas of relevance and priority area to a country, as well as the most available and accessible data. They provide an indication of the relative importance of genetic diversity for the country, which may also be focused on the available information and knowledge of the authors and may not be truly representative of actions on the ground.
